# High-throughput functional mapping of variants in an arrhythmia gene, *KCNE1*, reveals novel biology

**DOI:** 10.1101/2023.04.28.538612

**Authors:** Ayesha Muhammad, Maria E. Calandranis, Bian Li, Tao Yang, Daniel J. Blackwell, M. Lorena Harvey, Jeremy E. Smith, Ashli E. Chew, John A. Capra, Kenneth A. Matreyek, Douglas M. Fowler, Dan M. Roden, Andrew M. Glazer

## Abstract

**Background:** *KCNE1* encodes a 129-residue cardiac potassium channel (I_Ks_) subunit. KCNE1 variants are associated with long QT syndrome and atrial fibrillation. However, most variants have insufficient evidence of clinical consequences and thus limited clinical utility.

**Results:** Here, we demonstrate the power of variant effect mapping, which couples saturation mutagenesis with high-throughput sequencing, to ascertain the function of thousands of protein coding KCNE1 variants. We comprehensively assayed KCNE1 variant cell surface expression (2,554/2,709 possible single amino acid variants) and function (2,539 variants). We identified 470 loss-of-surface expression and 588 loss-of-function variants. Out of the 588 loss-of-function variants, only 155 had low cell surface expression. The latter half of the protein is dispensable for protein trafficking but essential for channel function. 22 of the 30 KCNE1 residues (73%) highly intolerant of variation were in predicted close contact with binding partners KCNQ1 or calmodulin. Our data were highly concordant with gold standard electrophysiological data (ρ = −0.65), population and patient cohorts (32/38 concordant variants), and computational metrics (ρ = −0.55). Our data provide moderate-strength evidence for the ACMG/AMP functional criteria for benign and pathogenic variants.

**Conclusions:** Comprehensive variant effect maps of *KCNE1* can both provide insight into I_Ks_ channel biology and help reclassify variants of uncertain significance.

## Background

Loss-of-function variants in *KCNE1* can cause type 5 long QT syndrome (LQT5; MIM 613695), a cardiac arrhythmia disorder characterized by QT prolongation on the electrocardiogram and an increased risk of sudden cardiac death.^1^ Heterozygous LQT5 variants can cause isolated QT prolongation (also known as Romano-Ward Syndrome),^2^ and homozygous and compound heterozygous variants can cause both QT prolongation and deafness (Jervell and Lange-Nielsen Syndrome; MIM 612347).^3^ Almost 50 unique *KCNE1* variants have been observed in patients with LQT5.^1, 4–6^ In addition, two gain-of-function variants have been associated with atrial fibrillation in isolated families.^7^ *KCNE1* encodes a single-spanning transmembrane protein that acts as a modulatory subunit to the pore-conducting KCNQ1 to form the I_Ks_ channel critical for cardiac repolarization (Figure 1A). The I_Ks_ complex is conventionally thought of as comprising KCNE1 and KCNQ1. However, additional proteins, including calmodulin, interact with the complex to modulate channel electrophysiology.^8^ *KCNE1* variants can disrupt channel function in two major ways: by reducing cell surface expression or by altering gating to reduce potassium flux.^9^

**Figure 1:**
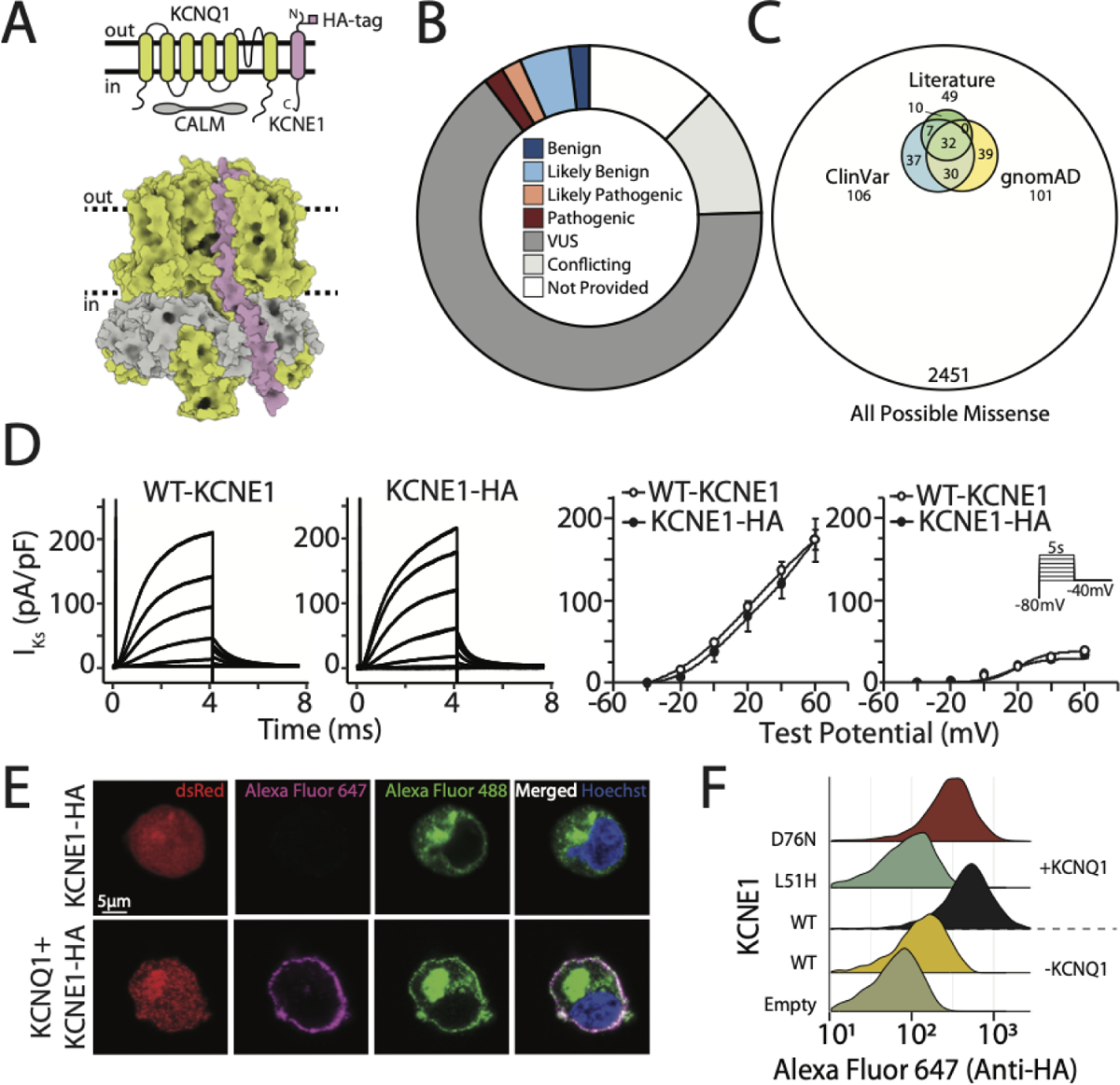
Overview of the I_Ks_ channel and a novel location for the HA epitope. A) The I_Ks_ channel complex is formed by KCNE1 (pink), KCNQ1 (green), and calmodulin (gray; top: schematic, and bottom: three-dimensional structure). The HA tag in KCNE1 (cloned between residues 34 and 35) is labeled. B) Most of the 106 missense KCNE1 variant classifications in the ClinVar database are VUS or conflicting (Figure style adapted from Starita et al., 2017). C) Variants reported in ClinVar, gnomAD, and previous studies comprise a small portion of all possible KCNE1 missense variants. D) The HA tag did not disrupt I_Ks_ (left: representative traces, inset: voltage protocol, and right: quantification of peak and tail current; n=4 cells/condition). E) An HA tag was used to visualize KCNE1. Live HEK293T cells were stained with an Alexa Fluor 647-conjugated anti-HA antibody to detect surface KCNE1, then fixed, permeabilized, and stained with an Alexa Fluor 488-conjugated anti-HA antibody to detect total KCNE1. dsRed is the KCNQ1 and KCNE1 transfection marker. Cells expressing only KCNE1-HA showed minimal surface labeling in the absence of KCNQ1 (top). Cells expressing KCNE1-HA and KCNQ1 had normal cell surface expression (bottom). F) Live cells stained for extracellular KCNE1-HA had minimal but detectable labeling in the absence of KCNQ1, but 6.1-fold higher labeling in the presence of KCNQ1. Known trafficking-null variant (L51H) had minimal cell surface expression, but a known gating variant (D76N) had near-wild type cell surface expression.

The American College of Medical Genetics and Genomics/Association of Molecular Pathology (ACMG/AMP) criteria are used to classify variants as pathogenic, likely pathogenic, likely benign, or benign.^10^ Variants with inconclusive evidence for clinical classification are designated as variants of uncertain significance (VUS). For many Mendelian disease genes, including *KCNE1*, most discovered variants are VUS and thus have limited impact on clinical decision-making,^10^ presenting a major challenge to genomic medicine.^11, 12^ In the ClinVar database of variant classifications,^13^ 77.4% of *KCNE1* variants are either VUS or have conflicting interpretations (Figure 1B). These variants, together with those reported by gnomAD or clinical case studies, represent a small subset of all possible missense variants (Figure 1C). It has been proposed that all missense variants compatible with life likely already exist in ∼50 individuals on the planet.^14^ In the ACMG/AMP scheme, well-validated *in vitro* functional studies can provide up to strong-level evidence for the PS3 (damaging effect on protein) and BS3 (no damaging effect) criteria.^15, 16^ However, functional assessment lags behind the rate of VUS discovery due to the rapid increase of genetic testing in research and clinical domains.^12^ One way to address this VUS problem is to leverage multiplexed assays for variant effects (MAVEs) to test thousands of variants in a single, highly-multiplexed experiment.^12, 17^ In a MAVE, a comprehensive variant library is coupled to a selection assay and high-throughput sequencing to ascertain variant function.

In this study, we conducted multiplexed assays of variant cell surface expression and potassium flux on a library of 2,592 single residue KCNE1 variants (95.7% of 2,709 total possible variants, Figure S1). We used an HA epitope in KCNE1 to determine cell surface expression of 2,554 individual variants using a “landing pad” cell line,^18^ which integrates a single variant per cell. This assay identified 470 missense variants that decrease and 310 that increase cell surface expression. We also used a gain-of-function KCNQ1 variant to design a cell survival assay that selects against cells with functioning channels. We deployed the function assay on 2,539 KCNE1 variants and found 588 loss-of-function missense variants, only 155 of which also had low cell surface expression. These datasets correlate well with previously validated *in vitro* studies, computational predictors, population metrics, and clinical phenotypes. Our data provide insight into KCNE1 structure and biology and can be implemented to reclassify variants in the ACMG/AMP scheme.

## Results

### Validation of an extracellular KCNE1 HA tag to detect cell surface expression

We cloned a 9-residue hemagglutinin (HA) tag into the extracellular domain of *KCNE1* (between residue 33 and 34; Figure 1A). KCNE1-HA (33-34) resulted in 3-fold higher cell surface labeling of the I_Ks_ channel complex compared to a previously described HA tag located in a predicted alpha helix (Figure S2A).^19^ Neither I peak and tail current densities, nor activation/inactivation properties were significantly different between KCNE1-WT and KCNE1-HA (Figure 1D). Human Embryonic Kidney (HEK) 293T landing pad cells expressing *KCNE1*-HA were labeled with a fluorophore-conjugated anti-HA antibody, visualized by confocal microscopy, and quantified by flow cytometry (Figures 1E and 1F). In cells expressing *KCNE1*-HA only, minimal plasma membrane anti-HA labeling was present, consistent with previous studies.^9, 20^ In contrast, cells co-expressing both *KCNE1*-HA and *KCNQ1* demonstrated substantial plasma membrane anti-HA labeling in live cells, and both plasma membrane and intracellular labeling in permeabilized cells (Figure 1E).^9, 21^ Quantification of cell surface expression by flow cytometry confirmed these observations (Figure 1F). A minimal but detectable amount of plasma membrane labeling was seen in *KCNE1*-HA only cells but the addition of *KCNQ1* caused a 6.1-fold increase in cell surface expression (Figures 1F and S2B). As expected, a known loss-of-trafficking variant (L51H) had minimal cell surface expression and a known gating variant (D76N) had near-WT cell surface expression when co-expressed with *KCNQ1* (Figures 1F and S2B).^22^

### A comprehensive library of KCNE1 mutations

We generated a comprehensive variant library of the 129-residue KCNE1 protein associated with random 18-mer barcodes using inverse PCR mutagenesis with degenerate primers (See *Methods*, Figure S3A).^23^ We linked 80,282 barcodes to 2,368 missense, 100 synonymous, and 124 nonsense KCNE1 variants (95.7% of the total possible variants; Figure S3B). Each variant was represented by a mean of 31 barcodes (Figure S3C).

### Multiplexed assay of KCNE1 cell surface expression

We coupled antibody labeling of the extracellular HA tag to deep sequencing of the library barcodes to perform a multiplexed assay of KCNE1 cell surface expression in cells engineered to constitutively express *KCNQ1* (Figure 2A). After quality control (see *Methods*; Figure S4A), we obtained cell surface trafficking scores for 98 synonymous, 117 nonsense, and 2,339 missense variants (94.2% of all possible variants; Figure 2B). Scores calculated from three individual replicates were highly concordant (Spearman’s ρ = 0.81-0.87, p < 0.001; Figure S4B). Cell surface trafficking scores for synonymous variants were normally distributed (0.988-1.024, 95% CI, Figures 2B-C). Nonsense variant scores were bimodally distributed (mode 1 = 0.01, mode 2 = 0.74). Nonsense variants from residue 1 to 55 were all trafficking-deficient, whereas nonsense variants at or after residue 56 had near-WT or higher than WT cell surface expression (p = 2.5×10^-20^, Wilcoxon test; Figures 2B-C). We therefore classified nonsense variants as early nonsense (residue 1-55) and late nonsense (56-129) in all subsequent trafficking analyses. All variants at residue 1 were trafficking-null as expected.

**Figure 2:**
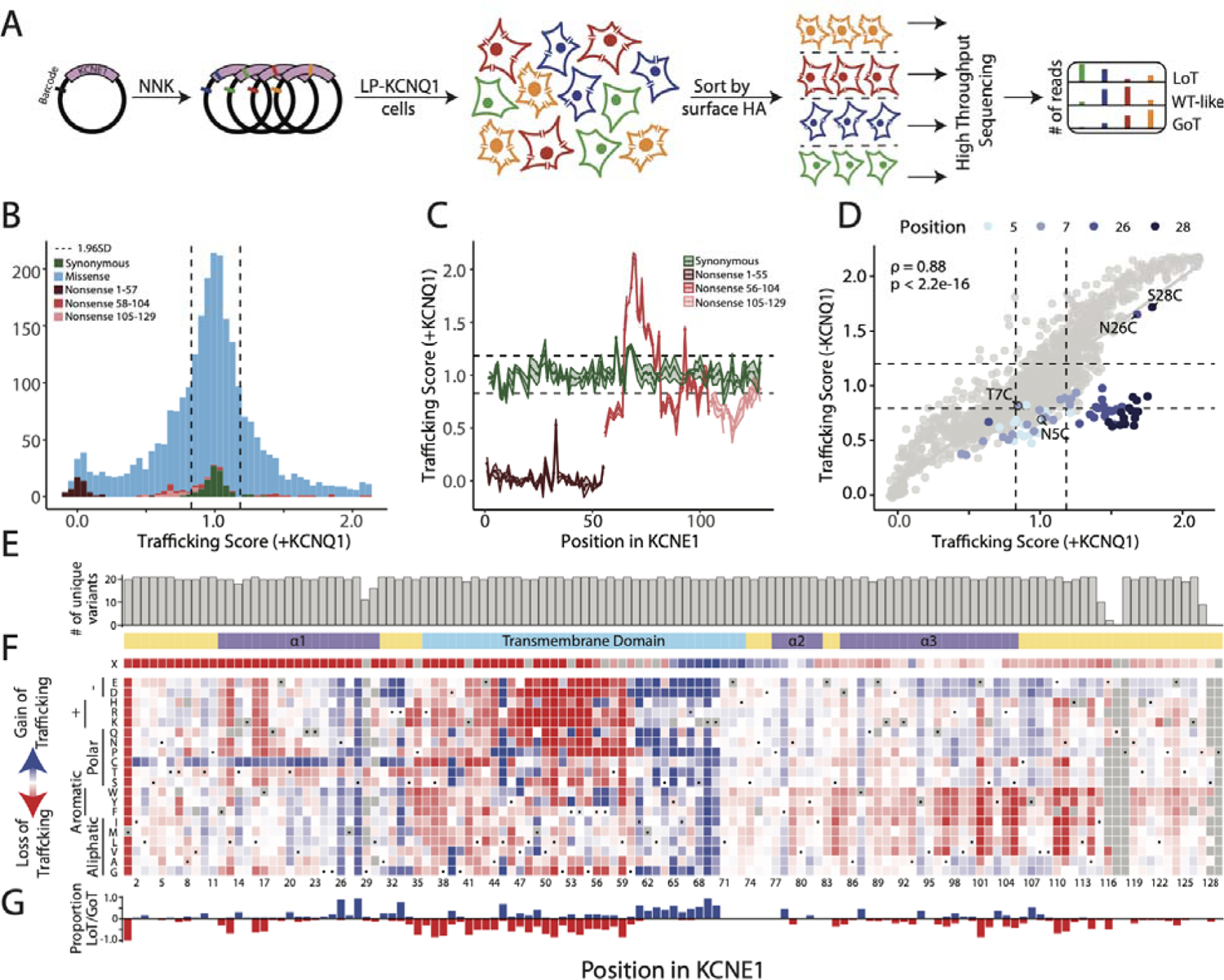
Multiplexed assay of KCNE1 cell surface expression. A) To conduct the cell surface expression multiplexed assay of KCNE1, a comprehensively mutated barcoded plasmid pool was integrated into the LP-KCNQ1 cell line. Cells were stained and sorted by surface KCNE1 levels and deep sequenced. B) Histogram of normalized trafficking scores by functional class showed a unimodal distribution of missense variant scores centered at the median of synonymous variant scores. A smaller peak at the median of early nonsense variant scores and a tail of super-trafficking variant scores was also seen (Dotted lines: mean ± 1.96 SD for the synonymous distribution). C) Most nonsense variants (1 per residue) from residues 1-55 (brown) are loss-of-trafficking but nonsense variants after residue 55 (red or pink) are gain-of-trafficking or WT-like. Synonymous variants (1 per residue) are represented in green (Dotted lines: mean ± 1.96 SD for the synonymous distribution). D) Trafficking scores in the presence and absence of KCNQ1 were highly correlated with the exception of variants at residues 5, 7, 26, and 28 which disrupted glycosylation sequons (highlight). Cysteine variants at these sequons did not follow this trend (annotation). See Figure S5 for a full description of the KCNQ1 trafficking assay (Dotted lines: mean ± 1.96 SD for the synonymous distribution). E) Near complete representation of variants was scored at each position in the KCNE1 protein (max = 21). F) A heatmap showing trafficking scores. Red, white, and blue indicate low, WT-like, and high cell surface expression, respectively. WT amino acids are indicated at each position with a dot. The colored ribbon indicates secondary structure, (purple = alpha helices, light blue = transmembrane domain, yellow = unstructured regions). G) For each residue, the proportion of gain-of-trafficking (GoT) and loss-of-trafficking (LoT) missense variants is displayed.

Missense variants with score estimates (95% confidence interval) lower than 2.5^th^ percentile or higher than 97.5^th^ percentile of the synonymous variant distribution were assigned as loss- or gain-of-trafficking respectively. Using these cutoffs, we identified 470 loss-of-trafficking variants and 311 gain-of-trafficking variants. We also compared trafficking scores to the few literature reports of cell surface trafficking measurements, typically reported as “normal” or “loss-of-trafficking.” 7/9 variants were concordant: 6 variants with normal or near-normal trafficking (T6F, S74L D76A, D76N, Y81C, and W87R) and 1 loss-of-trafficking variant (L51H; Figure S4C).^22, 24–26^ Two discordant variants, R98W and G52R, were previously reported to have normal trafficking but were loss-of-trafficking in our assay.^9^

Several interesting and previously unreported findings emerged from the cell surface trafficking scores (Figure 2F). As mentioned above, nonsense variants after residue 55 were still able to traffic to the cell surface. In the extracellular N-terminus, almost all cysteine variants from residue 2 to 33 had increased cell surface expression. Residues 46-59 in the transmembrane helix, likely juxtaposed to the hydrophobic core of the lipid bilayer, were highly resistant to polar or charged variation. Variants in residues 61-70, interacting with the intracellular hydrophilic edge of the cell membrane, increased cell surface expression. Predicted alpha helices in the intracellular C-terminus (residues 79-114) had multiple residues resistant to aromatic or larger aliphatic variation.

As HEK293T cells only expressing KCNE1 exhibited detectable levels of anti-HA staining, we conducted a second MAVE of KCNE1 cell surface expression in the absence of co-expressed KCNQ1 (“KCNQ1-”; Figure S5) to explore their intermolecular interaction. Trafficking scores for missense variants in the presence and absence of *KCNQ1* were highly correlated (Figure 2D, Spearman’s ρ = 0.88, p < 2×10^-6^), except for variants in the N-glycosylation sequons (NXS/NXT motifs) at positions 5-7 and 26-28 in the extracellular N-terminus of KCNE1.^26, 27^ Most variants disrupting N-glycosylation especially at residues 26 and 28 reduced cell surface expression in the absence of KCNQ1 (Figure 2D). Thus, loss of N-terminal glycosylation is more advantageous to cell surface expression in the presence of KCNQ1. Cell surface expression of cysteine super-trafficking variants (including at the 5-7 and 26-28 glycosylation sequons) was not affected by KCNQ1. As discussed below, this result may reflect antagonism between KCNE1 N-glycosylation and KCNE1/KCNQx interaction unique to KCNQ1 and not other binding partners possibly expressed in HEK293T cells.

### Development of a cell fitness assay for KCNE1 function

Since I_Ks_ channels are closed at the resting potential of HEK293T cells (approximately −25 mV),^28^ we developed an assay for KCNE1 function by leveraging the electrophysiology of a previously described gain-of-function variant, KCNQ1-S140G.^29, 30^ In the presence of KCNE1, KCNQ1-S140G left-shifts the voltage of I channel activation^29^ and reduces channel deactivation;^31^ in the absence of KCNE1, the channel expresses minimal current (Figure 3A). These electrophysiological shifts result in substantial KCNE1-dependent K^+^ efflux at −25 mV. We hypothesized that long-term expression of I_Ks_ channels formed by KCNQ1-S140G and normally functioning KCNE1 would decrease cell fitness due to continual K^+^ efflux. To test this hypothesis, we integrated a 1:1 mixture of empty vector control and either wild-type KCNE1-HA or previously studied KCNE1 variants into HEK293T landing pad cells stably expressing KCNQ1-S140G (LP-KCNQ1-S140G, Figure 3B). We then quantified KCNE1^+^ and KCNE1^-^ cell proportions over 20 days (Figure 3C and 3D). Compared to cells with empty vector controls, cells with WT KCNE1 depleted over time, whereas cells with loss-of-function KCNE1 variants (L51H or D76N) persistently survived. Two previously described KCNE1 gain-of-function variants (G60D and G25V)^7^ depleted at the same rate as KCNE1-HA-WT (Figure 3D), indicating that this assay does not distinguish gain-of-function variants from WT.

**Figure 3:**
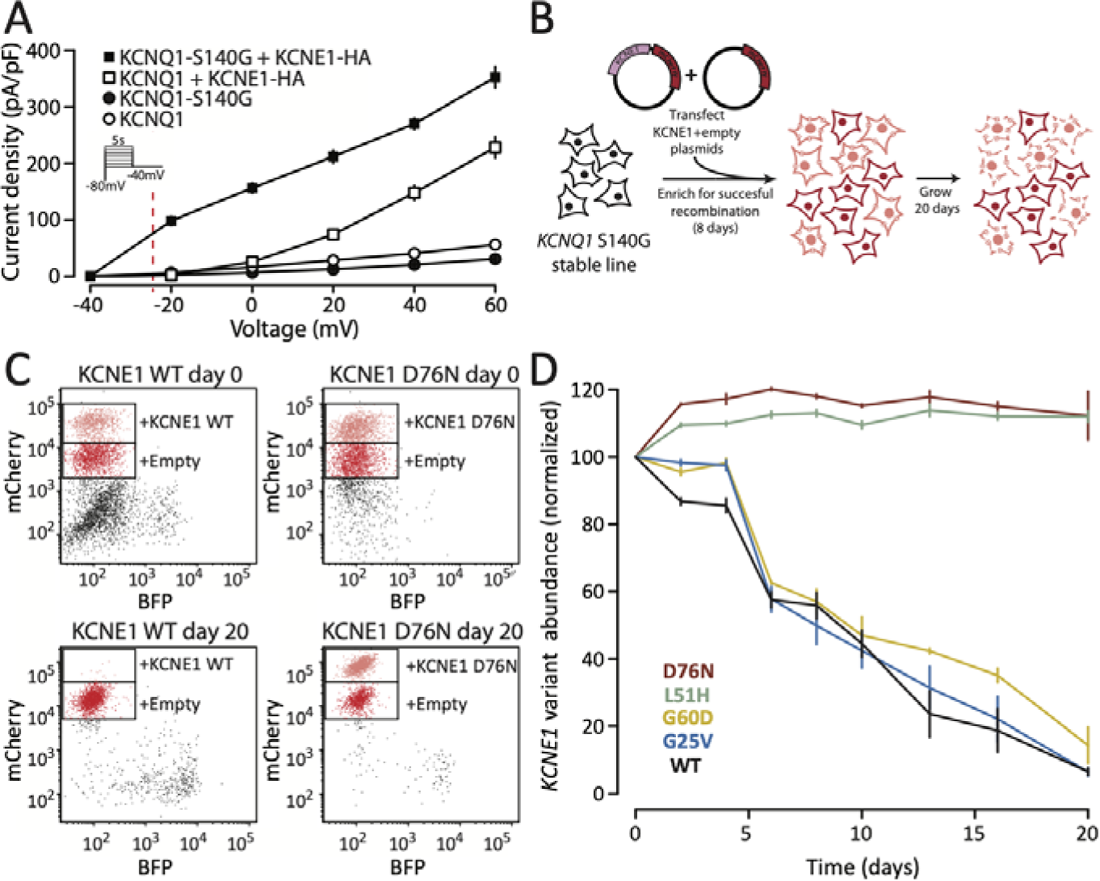
A KCNE1 function assay using a gain-of-function KCNQ1 variant. A) Voltage clamp measurements of outward potassium currents in HEK293T cells transfected with various combinations of WT or S140G KCNQ1 and/or KCNE1-HA. At the resting potential of HEK293T cells (∼ −25 mV, dotted line), there is almost no potassium current in cells transfected with KCNQ1 ± KCNE1-HA. However, the gain-of-function KCNQ1-S140G variant results in a leftward shift of voltage of activation and increased current at −25 mV, only in the presence of KCNE1. B) Experiment to validate KCNE1 selection assay using LP-KCNQ1-S140G cells transfected with 1:1 pools of KCNE1 variant and empty vector plasmids. C) Representative flow cytometry measurements of cells transfected with 1:1 pools of plasmids. Cells with empty vectors had reduced mCherry fluorescence allowing quantification of distinct cell populations. After 20 days of selection, there was a strong depletion of cells expressing KCNE1-HA-WT but not a loss-of-function variant D76N. D) Time course of relative fitness of cells expressing KCNE1 variants compared to cells expressing empty vectors. All values are normalized to the values at day 0. Cells expressing KCNE1-HA-WT or two gain-of-function variants (G25V and G60D) depleted over 20 days but cells expressing two loss-of-function variants (D76N and L51H) persisted in the cell pool. Points and error bars indicate mean and standard error of three replicates.

### Multiplexed assay of KCNE1 function

We used this KCNQ1-S140G-based cell fitness assay to conduct a multiplexed assay of KCNE1 function. We integrated the KCNE1 library into HEK293T cells stably expressing KCNQ1-S140G and quantified cell survival over 20 days (Figure 4A). After 20 days, cell expressing synonymous variants were depleted and those expressing most nonsense variants persistently survived (Figure S6A). These changes in the cell pool were not seen in the presence of WT-KCNQ1 (Figure S6B). Variant depletion from day 0 to day 20 was used to calculate normalized KCNE1 function scores for 98 synonymous, 121 nonsense, and 2,320 missense variants (93.7% of all variants; Figures 4B and S6C). Function scores across three replicates were highly concordant (Spearman’s ρ=0.87; Figure S6D). Synonymous variants followed an asymmetric unimodal distribution (median = 0.99, IQR=0.27, Figure 4B). Nonsense variants were bimodally distributed (mode 1 = 0.02, mode 2 = 0.99); nonsense variants at residues 1-104 were loss-of-function, whereas nonsense variants at residue 105-129 had WT-like function (p = 2.6×10^-13^, Wilcoxon test, Figure 4C). Thus, across the trafficking and function assays, three classes of nonsense variants were present based on residue number: #1-55 caused loss of trafficking and function, #56-104 caused WT-like or elevated trafficking but loss-of-function, and #105-129 caused near WT-like trafficking and function (Figure S7A). All variants at residue 1 were loss-of-function as expected. Missense variant function scores were bimodally distributed (mode 1 = 0.14, mode 2 = 1.04). The two modes approximately corresponded to the peaks of the synonymous distribution and early nonsense distributions. Upon comparing trafficking and function scores for missense variants, we found 136 trafficking-null and 288 trafficking-normal, functionally deleterious variants (Figure 4D). We refer to the latter as “gating variants.”

**Figure 4:**
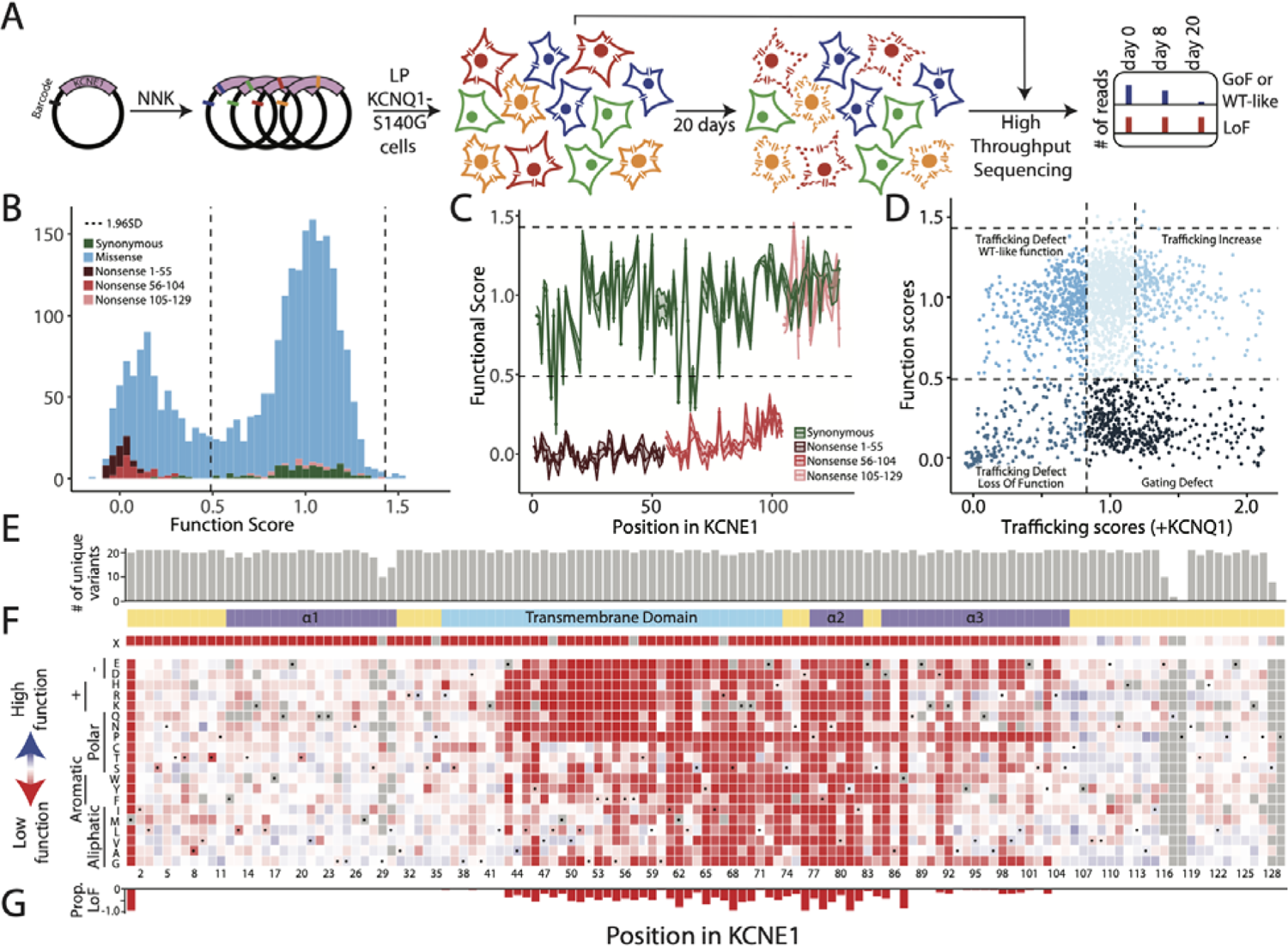
Multiplexed assay of KCNE1 function. A) Schematic of multiplexed assay of KCNE1 function. A comprehensively mutated barcoded plasmid pool was integrated into LP-KCNQ1-S140G cells. Cells were grown for 20 days and cell pools at 0 days, 8 days, and 20 days were deep sequenced. B) Histogram of normalized functional scores by functional class showed a bimodal distribution of missense variant (Dotted lines: mean±1.96 SD for the synonymous distribution). C) Nonsense variants (1 per residue) up to residue 104 are functionally deleterious (brown and red), including residues 56-104 that were dispensable for cell surface trafficking (red). Synonymous variants (1 per residue) are represented in green (Dotted lines: mean ± 1.96 SD for the synonymous distribution). D) Relationship between trafficking and function scores for missense variants showed that most deleterious variants disrupt gating (Dotted lines: mean ± 1.96 SD for the synonymous distribution). E) Near complete representation of missense variants was scored at each position in KCNE1 (max = 21). F) KCNE1 function score heatmap. Red, white, and blue indicate loss-of-function, normal function, and gain-of-function, respectively. WT amino acids are indicated at each position with a dot. The colored ribbon indicates secondary structure as in Figure 2. G) For each residue, the proportion of loss-of-function missense variants is displayed.

On the heatmap of KCNE1 function scores, variation-resistant regions were located in the transmembrane helix (residues 61-73) and the transmembrane-proximal cytosolic alpha helix (residues 77-82; Figure 4F). Known trafficking-null variants (L51H) and gating variants (D76N, W87R) had deleterious function scores.^22^ Common polymorphisms (G38S, D85N) had WT-like trafficking and function scores, consistent with their minimal effects on baseline QT interval. Residue 85 is otherwise resistant to variation and D85N is only one of 2 missense variants at this residue with WT-like score estimates (D85N: 0.79-1.02, and D85E: 0.69-0.98, 95% confidence interval). Variants that disrupted glycosylation sequons (residues 5-7 and 26-28) largely did not alter channel function despite their effects on KCNE1 trafficking (Figure S7B), consistent with previous work.^26, 27^ Additionally, N-terminal cysteine residues, while gain-of-trafficking, did not drastically affect function. This is expected, as the assay is not well-powered to detect gain-of-function variants. Only one variant (Y107R) met the “gain-of-function” criteria (score estimate > 97.5% percentile of synonymous scores) and was validated by manual patch clamp (Figure S7C).^26, 27^

### Structural model of the KCNE1/KCNQ1/calmodulin complex

An experimental structure of the KCNE1/KCNQ1/calmodulin complex has not been determined, so we examined multiple AI-based models (Figure S8). Aligning predicted structures of KCNE1 to a reported cryo-EM structure of the KCNQ1-KCNE3 complex indicated that the conformation obtained from AlphaFold-multimer modeling (See Supplemental Methods) was likely the most biologically meaningful (Figures 1A and 5). We therefore overlaid mean trafficking and functional scores on the AlphaFold-multimer structural model of the KCNE1-KCNQ1-calmodulin complex (Figure 5A-D). As discussed above, the hydrophobic core of the transmembrane helix (residues 44 to 60) did not tolerate polar or charged variation in both trafficking and function maps. Multiple variants in the cytosolic membrane-proximal region of the transmembrane domain (residues 61-70) increased cell surface expression but were functionally intolerant to variation. This region contains contacts with KCNQ1, including KCNE1 residues M62, Y65, and S68 (Figure 5F). Thus, variation in this region likely disrupts normal gating properties of the I_Ks_ channel. Polar and charged variation in an additional predicted intracellular helix (residues 85-105 including functionally constrained residues F78, Y81, I82, and W87) preserved cell surface expression. However, the helix was functionally intolerant to variation likely due to observed interaction with calmodulin (Figure 5G) that affects gating properties of the I_Ks_ complex. Overall, of the 30 KCNE1 residues highly intolerant of variation (>70% of missense variants at given residue position are loss-of-function), 22 (73%) were in close contact (<5Å) with KCNQ1 (14 residues) and/or calmodulin (9 residues; residue 77 was in close contact with both proteins). These variation-resistant residues comprise 40% of 55 KCNE1 residues in close contact with KCNQ1 and/or calmodulin. On the other hand, none of the 70 residues where >50% of the missense variants had WT-like functional scores were in close contact with either KCNQ1 or calmodulin.

**Figure 5:**
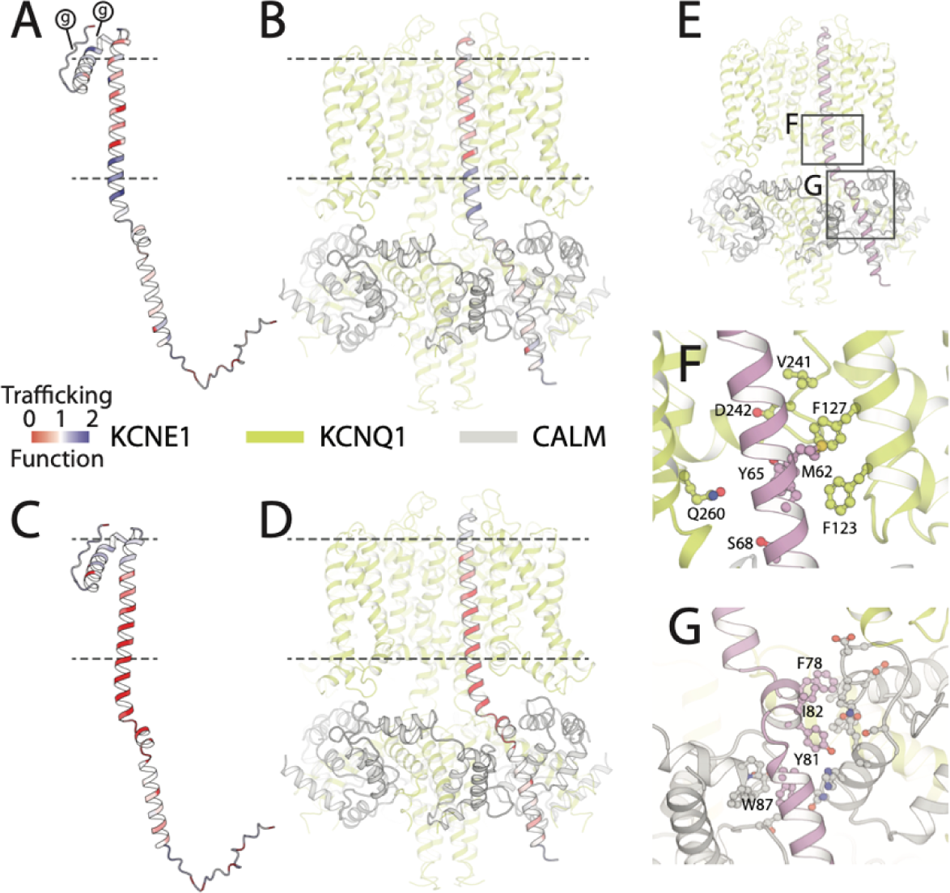
Structural impact of variant effects. A-D) KCNE1 structural model, coded by mean trafficking scores (A,B) or function scores (C,D) at each residue on the same color scale as in Figures 2F and 4F (Red = loss-of-trafficking/function, white = WT-like, blue = gain-of-trafficking/function). Dotted lines indicate the approximate location of the plasma membrane. (g) represents glycosylation sites. Panels B and D show KCNE1 in complex with KCNQ1 (green) and calmodulin (gray). E) I_Ks_ complex showing location of panels F and G. F) Constrained KCNE1 residues (pink) that make contacts with KCNQ1. KCNE1 residue M62 forms a hydrophobic cluster with KCNQ1 residues F123, F127 and V241, whereas KCNE1 residues Y65 and S68 make polar contacts with KCNQ1 residues D242 and Q260, respectively. G) Constrained KCNE1 residues (pink; F78, Y81, I82, W87) that make contacts with calmodulin. Residues on calmodulin not labeled.

### Correlation of MAVE scores with patch clamp and patient data

We curated a list of 149 *KCNE1* variants with previously published patch clamp, trafficking and patient data, (Additional File 1) and conducted additional patch clamp studies of 15 variants (Table S1). The MAVE function scores were strongly correlated with measured or reported peak currents (p = 1.5×10^-9^, ρ = 0.65, n = 71; Figure 6A). 26/29 variants with ≤25% of WT peak current had deleterious function scores. Only 8 of these 26 had deleterious trafficking scores (30.8%; Figure S9A). 21/29 variants with >50% WT peak current had WT-like function and trafficking scores. Although function scores correlated best with peak current, *KCNE1* variants with large shifts in the voltage or time kinetics of activation or deactivation were also more likely to have deleterious scores (Figures 6B and S9B). The trafficking scores had weaker correlations with patch clamp parameters (Figures S9C and S9D). Single nucleotide variants (SNVs) achieved by transversion mutations were more likely to be functionally deleterious compared to variants achieved by transition mutations (p = 2.2×10^-4^, Wilcoxon test; Figures S9E and S9F).

**Figure 6:**
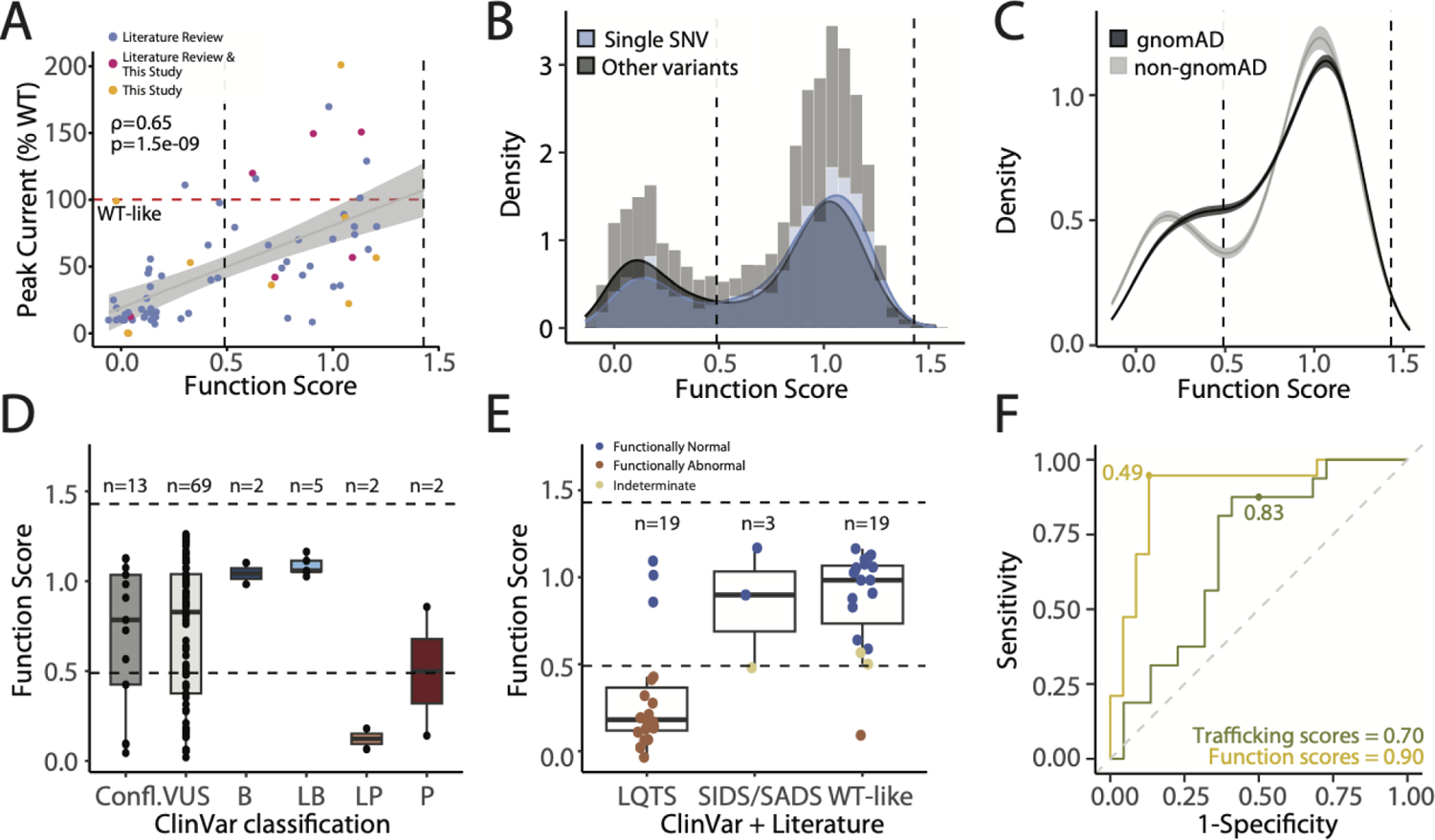
Function scores correlate with in vitro assays and clinical outcomes. A) Correlation of function scores with peak current (normalized to WT) from patch clamp studies. Blue: literature review, pink: literature review and this study, yellow: this study. B) Single SNV variants (blue) are less likely to have deleterious function scores compared to all other variants (black; p = 9.3×10^-6^, Wilcoxon Test). C) Variants present in gnomad (black) were more likely to have WT-like scores than variants absent from gnomAD (grey; p=2.2×10^-4^, Wilcoxon Test). Mean and confidence intervals are generated by re-sampling both distributions 100 times. D) Function scores by ClinVar classification. E) Function scores for presumed control and presumed disease-associated variants. See Additional File 1 for the literature review dataset. F) Receiver operator characteristic curves evaluating prediction of variant pathogenicity by function and trafficking scores. Dots indicate the deleteriousness cutoffs used in this study (functional scores = 0.49, trafficking scores = 0.83). The area under the curve is listed at the bottom of the panel.

We examined the distribution of missense variant scores in presumed normal and pathogenic variants, curated from gnomAD, ClinVar and our literature review of LQT5 and JLNS cases. Variants achieved by more than one single SNV or not present in gnomAD tended to have deleterious function scores (p= 9.3×10^-6^, Wilcoxon test in Figure 6B; p = 4.6×10^-5^, Wilcoxon test in Figure 6C; and Figure S10A). Function scores were largely concordant with clinical outcomes: 7/7 ClinVar benign/likely benign variants and 16/19 presumed normal variants had WT-like function scores; and 3/4 ClinVar pathogenic/likely pathogenic variants and 16/19 presumed pathogenic variants had deleterious function scores (Figures 6D and 6E). The predictive performance of MAVE scores was tested by generating receiver operator characteristic (ROC) curves. The area under the ROC curve was 0.90 for function scores and 0.70 for trafficking scores (Figure 6F). Function scores correlated well with several previously validated computational and evolutionary metrics of variant effect (Figure S11). The predictive performance of function scores was comparable to these metrics (Figure S10B). We note that while computational metrics perform well for *KCNE1* variants, our MAVE data are derived from direct *in vitro* data, and thus provide orthogonal information. These trends were weaker for trafficking scores (Figures S12 and S13).

We calibrated our assay with presumed pathogenic and benign variants, following the approach recommended by the ClinGen Sequence Variant Interpretation working group.^16^ We used the function scores for calibration because of their better correlation with clinical outcomes and other metrics of disease risk. Using the ClinVar only dataset, the functional assay achieved a likelihood ratio of pathogenicity (termed “OddsPath”) of 13.0 for pathogenic (PS3_moderate) and 0.253 for benign (BS3_supporting). Using the expanded set of presumed benign and pathogenic variants, the functional assay achieved an OddsPath of 14.3 for pathogenic (PS3_moderate) and 0.168 for benign (BS3_moderate).

## Discussion

A majority of variants in clinically relevant genes, including *KCNE1*, are VUS, and resolving the clinical significance of VUS at scale has been a major challenge in genomic medicine.^11^ High-throughput functional data can be deployed to reclassify as many as 89% of known VUS in clinically actionable genes.^15^ Loss-of-function *KCNE1* variants cause type 5 long QT syndrome (LQT5) via two major mechanisms: reduced cell surface expression and/or defective gating (activation of the I_Ks_ complex).^32^ Our study ascertains trafficking and electrophysiology properties to better understand the disease risk of nearly all possible *KCNE1* variants. Furthermore, our dataset provides functional scores for 871 of the 924 *KCNE1* variants accessible by a single SNV; each variant, if compatible with life, is estimated to exist in approximately 50 humans currently alive.^14^ Our comprehensive datasets also reveal new biology not apparent from previous smaller-scale mutational studies.

### Insights from trafficking scores

We identified 470 loss-of-trafficking and 310 gain-of-trafficking missense variants, including a stretch of super-trafficking N-terminal cysteine variants. We hypothesize that these residues may increase cell surface expression by forming intermolecular disulfide bonds with other proteins (e.g., KCNE1 or KCNQ1). We tested this hypothesis by conducting a MAVE of cell surface expression in the absence of KCNQ1, but this dataset may be influenced by other low-abundance binding partners of KCNE1 endogenous to HEK293T cells. Nevertheless, the result clearly implicated loss of N-glycosylation at position 26, but less so at position 5, as increasing cell surface expression in the presence of KCNQ1. The cell surface abundance of nonsense variants in our data also highlights that the latter half of the protein (residue 56-129) is dispensable for cell surface trafficking. This residue marks the beginning of the FTL domain, important for electrophysiological modulation of the I channel,^33, 34^ and corresponds to a previously reported kink in the transmembrane domain.^35, 36^

### Insights from function scores

We developed a novel selection assay that used a gain-of-function KCNQ1 variant (S140G) to identify loss-of-function KCNE1 variants. With the assay, we identified 588 loss-of-function variants, of which 433 (73.7%) had normal cell surface expression. We refer to these as “gating” variants, as proper K^+^ flux through the I_Ks_ channel is likely disrupted. The predominance of pathogenic gating variants is in contrast to other potassium channel disease genes, such as *KCNH2*; 88% of 193 studied KCNH2 loss-of-function variants disrupt cell surface expression.^37^ The high proportion of gating variants is likely explained by KCNE1’s primary function as a KCNQ1-modifying subunit to modulate the I_Ks_ complex. The gating variants identified in our data include well-studied variants like D76N and W87R^22^ and novel variants in the transmembrane-proximal cytosolic domain that interact with KCNQ1 and calmodulin. We also identified gating variants predicted to interact with KCNQ1 that otherwise increase KCNE1 cell surface expression. These variants likely do not disrupt intermolecular binding within the I_Ks_ channel complex but are rather involved in the physical modulation of KCNQ1 with appropriate electrochemical stimuli.

Our work defines three classes of nonsense variants based on residue location: #1-55 that disrupt cell surface expression, #56-104 traffic to the cell surface but do not function, and #105-128 have near-normal trafficking and function.^38^ While D85N (rs1805128; AF 0.14%-2.54% in gnomAD) is associated with QT prolongation and fatal arrhythmias in the presence of drugs and other environmental factors,^39^ the variant is too common to cause long QT syndrome in isolation. The WT-like MAVE score estimates are concordant with this observation (trafficking score: 0.82-1.00; functional score: 0.79-1.02).

### Structural implications of MAVE data

Although there is no solved cryo-EM structure of the KCNQ1-KCNE1-calmodulin complex, we used AlphaFold2 and homology modeling to construct a structural model of the I_Ks_ complex. In our MAVE datasets, hydrophilic variants in the hydrophobic transmembrane core (including previously studied variants like L51H)^22^ disrupt function by affecting cell surface expression. We also identify a 26 amino acid membrane-proximal cytosolic region (residues 61-87) that comprises essential elements for I_Ks_ complex function: KCNQ1-binding and calmodulin-binding regions, and a structured linker connecting the two. Variants in this region likely disrupt hydrophobic and polar interactions and pi-stacking with the corresponding residues of KCNQ1 and calmodulin respectively. Accordingly, the region is highly constrained in the function, but not trafficking, assay. With increasing confidence in and accuracy of AI-generated structural models, the resolution and interpretation of MAVE-based structure-function relationships will likely improve.

### Clinical Associations of deleterious KCNE1 variants

Our dataset also highlights the strength of association between *KCNE1* loss-of-function variants and QT prolongation. In 2020, the ClinGen Channelopathy Working Group assessed *KCNE1* as having only limited evidence for association with congenital type 5 long QT syndrome (LQT5) and strong evidence for association with acquired long QT syndrome.^40^ Since then, several papers have further described the LQT5 risk of *KCNE1* variants. Roberts *et al*.^1^ reported in a multi-center review of LQT5 that *KCNE1* variants are a low penetrance cause of congenital long QT syndrome. Our group has previously shown that carriers of pathogenic/likely pathogenic *KCNE1* variants in the electronic MEdical Records and GEnomics (eMERGE) sequencing study had longer QT intervals and higher odds of arrhythmia diagnoses and phenotypes.^41^ However, these variants conferred a lower risk than P/LP variants in *KCNQ1* or *KCNH2* (LQT1 and LQT2). The QT prolonging effects of deleterious *KCNE1* variants has also been established in both the TOPMed and UK Biobank cohorts, with effect sizes up to 15-20 ms.^42^ Thus, *KCNE1* loss-of-function variants appear to contribute to modest increases in baseline QT interval.

The presence of a KCNE1 deleterious variant may not be sufficient for development of the congenital LQT5 phenotype. For the QT interval to rise above a high-risk threshold, LQT5-associated variant carriers may require additional genetic or environmental insults, such drug exposure.^1^ Based on the strong correlation between MAVE scores and functional and clinical outcomes, variants identified as functionally deleterious in this study likely predispose carriers to LQT5.

### MAVE data can supplement ACMG criteria

Both our trafficking and function assays had excellent separation between early nonsense and synonymous variants, high concordance among replicates, and strong correlations with previous measurements of cell surface trafficking or patch clamp function. In addition, the function MAVE dataset correlates well with patient and population cohorts, using controls curated from ClinVar, literature reports, and gnomAD. Another advantage of the MAVE approach is the uniformity and replication of the platform used to analyze variants, unlike literature reports derived from multiple individual laboratories with low concordance.^43^ Because of this heterogeneity, data from literature reports cannot accurately contribute to ClinGen guidelines for interpreting *in vitro* functional datasets.^16^ The small number of discordant variants between our assay and the literature might represent a failure of our assay or of previous reports. Based on the ClinGen guidelines, function scores can be implemented at the moderate level for both PS3 and BS3 criteria to aid in variant interpretation. However, given the higher clinical burden of false negatives, we recommend using our scores conservatively to apply BS3 at the supporting level. This work adds to the growing set of arrhythmia genes previously assayed by multiplexed assays for either the full protein (KCNJ2, calmodulin)^44, 45^ or a portion of the protein (SCN5A, KCNH2).^32, 46, 47^

## Limitations

The MAVE function assay was not well-powered to detect gain-of-function variants. The scores were generated in a heterologous system that may not fully model the behavior of the I_Ks_ channel in a cardiomyocyte. Additionally, we did not model several known behaviors of the I_Ks_ channel, such as regulation by PIP_2_ or adrenergic stimulation. We also did not measure dominant negative properties of the channel, demonstrated for some KCNE1 variants including D76N.^48^ HEK293T cells endogenously express non-KCNQ1 binding partners of KCNE1 that might confound trends when assessing the trafficking of KCNE1, especially in the absence of KCNQ1.

While variants with the lowest trafficking scores uniformly had loss-of-function scores, some partial loss-of-trafficking variants had non-deleterious function scores (Figure 4D). This result may reflect the flexible stoichiometry of the channel. KCNE1 can bind to KCNQ1 in a 1:4 to 4:4 conformation.^49^ Since KCNE1 is highly expressed in our assays, cells expressing partial loss-of-trafficking variants may bind to KCNQ1 in higher proportions to activate KCNQ1 and yield some I_Ks_ function.

Although KCNE1’s major role in the cardiac action potential is through its role in I_Ks_, KCNE1 has also been reported to interact with other proteins including KCNH2^50^ and TMEM16A, a cardiac chloride channel.^51^ These interactions might influence arrhythmia risk that is not captured by the I_Ks_ assay performed in this study. For example, previously discussed variant D85N has been reported to cause a partial loss-of-function of KCNH2,^52^ which may explain its influence on the QT interval. An extension of this work would be to investigate the influence of comprehensive KCNE1 variant libraries on other protein complexes.

## Conclusions

We comprehensively ascertained variant function in an important arrhythmia gene, *KCNE1*. We identified 470 variants that affect KCNE1 trafficking and 588 variants that alter function. Our work highlights new biology of the I_Ks_ channel and can be implemented in the ACMG/AMP scheme to classify variants.

## Declarations

### Ethics approval and consent to participate

Not applicable.

## Consent for publication

Not applicable.

### Availability of data and materials

The three MAVE datasets supporting the conclusions of this article are available in Additional File 2 and will be deposited in MaveDB^53^ upon publication. Illumina sequence data will be archived in the Short Read Archive upon publication. Code to analyze MAVE data and re-create figures is available at https://github.com/GlazerLab/KCNE1_DMS.

## Author Contributions

AM, DMR, and AG designed the research and wrote the manuscript. AM, MC, BL, TY, DJB, MLH, JS, AEC, and AMG performed the research. AM, MC, BL, TY, DJB, JS, DMR, and AMG analyzed the data. JAC, KAM, and DMF contributed essential resources and reagents. All authors reviewed and approved the manuscript.

## Acknowledgements

We thank Fritz Roth, Jochen Weile, Bjorn Knollmann, David Samuels, the CardioVar consortium, and members of the Vanderbilt Center for Arrhythmia and Therapeutics for helpful discussions. We thank Kara Skorge, Laura Short, Marcia Blair, and Lynn Hall for cloning assistance and the Vanderbilt VANTAGE sequencing core for performing Illumina sequencing.

## Author’s Twitter handles

@ayeshamyar (AM), @capra_lab (JAC), @kmatreyek (KAM), @dougfowler42 (DMF), @rodendm (DMR), and @amglazer (AMG)

## Methods

### Overview of multiplexed assays of KCNE1 variant effects

An HA epitope tag was cloned into the extracellular region of *KCNE1*. Saturation mutagenesis was used to create a plasmid library of 2,592 distinct single amino acid *KCNE1* variants (95.7% of total possible variants). Each plasmid was associated with a random 18-base barcode and the plasmid library was “subassembled”, i.e., deep sequenced to connect each barcode to its corresponding KCNE1 variant. To express one *KCNE1*-HA variant per cell, the library was integrated into Human Embryonic Kidney (HEK) 293T cells engineered to include a “landing pad.”^18^ To determine cell surface expression, *KCNE1*-HA variants were expressed with or without WT *KCNQ1* and the cell pool was stained with an anti-HA antibody. Cells were sorted into 4 bins based on the level of cell surface labeling, and each bin was deep sequenced to quantify variants. To determine channel function, *KCNE1-HA* variants were expressed with a gain-of-function *KCNQ1* variant for 20 days to deplete cells with normally functioning I_Ks_ channels. The cell pool was deep sequenced at day 0, 8 and 20 to quantify variants (Figure S1).

### Cloning a novel KCNE1 HA tag

We used *KCNE1* (NCBI Reference Sequence NM_001127670.4) with the G allele at residue 38 (p.S38G, c.112A>G, rs1805127), as this variant allele is more common globally (65.8% allele frequency [AF] across all ancestral populations in gnomAD; range 57.4%-71.3%).^54^ An HA epitope (5’-TACCCCTACGATGTACCAGATTATGCG-3’) was cloned between residues 34 and 35 with primers ag424 and ag425 using inverse PCR with Q5 polymerase (New England Biolabs, NEB).^23^ The sequences of all primers used in this study are presented in Table S2. The resulting gene, *KCNE1*-HA, was subcloned into the pIRES2-dsRED2 plasmid using PCR with ag409 and ag410 and NotI restriction digestion (NEB) to create *KCNE1-*HA:pIRES2:dsRED2. All plasmid sequences were verified with Sanger sequencing. Using NotI restriction digestion, the *KCNE1*-HA gene was moved to an AttB plasmid containing pIRES:mCherry:BlastR to create p*KCNE1*-HA:IRES:mCherry:BlastR. We also used a compact (<4kb) promoterless plasmid for single-variant mutagenesis.^46^ Maps of key plasmids used in this study are shown in Figure S14A.

### Generation of HEK293T lines stably expressing KCNQ1 or KCNQ1*-S140G*

The experiments were conducted in either previously published Bxb1-mediated landing pad HEK293T (LP) cells,^18^ or LP cells genetically engineered to express *KCNQ1* (WT or S140G). LP cells contain an AttP integration “landing pad,” Blue Fluorescent Protein (BFP), and iCasp9 caspase downstream of a doxycycline-induced promoter.^18^ The AttB:mCherry:BlastR plasmid integrates into the AttP landing pad. Prior to integration, the promoter drives BFP and iCasp9 expression. After integration, the promoter drives expression of mCherry and blasticidin resistance genes instead. Cells with a successful plasmid integration event are blasticidin resistant, express mCherry and lack iCasp9, which can be used to select for recombined cells. Genetically engineered LP cells expressing KCNQ1 (WT or S140G) were generated as below.

Since the plasmids use NotI restriction digestion for subcloning, we removed an intra-cDNA NotI restriction enzyme site, by introducing a synonymous mutation (c.246G>T), into WT *KCNQ1* cDNA (NM_000218.3) using QuikChange mutagenesis (Agilent). The *KCNQ1* gain-of-function variant (p.S140G, c.418A>G, rs120074192) was also generated by QuikChange mutagenesis. KCNQ1 S140G biases I_Ks_ channels to the open state^29, 31^ at lower membrane potentials (including the resting potential of HEK293T cells, approximately −25 mV)^28^ and leads to increased cellular K^+^ efflux.^55^

To construct a cell line with constitutive *KCNQ1* expression, *KCNQ1* (WT or p.S140G) was cloned into a Sleeping Beauty transposon system, pSBbi-GN (a gift from Eric Kowarz, Addgene# 60517),^56^ using SfiI restriction digestion (NEB) to create pSBbi-GN::KCNQ1. This plasmid expresses *KCNQ1* and *EGFP* via constitutively active bidirectional promoters. The plasmid was randomly integrated into a non-landing pad locus in the LP cells with the Sleeping Beauty transposase as follows. Cells were cultured at 37° at 5% CO_2_ in HEK media: 10% FBS (Corning), 1% non-essential amino acids (Corning), 1% penicillin/streptomycin (Corning) in DMEM (Thermo). On day 0, pSBbi-GN::KCNQ1 was transfected into the LP cell line along with a plasmid expressing the Sleeping Beauty transposase, pCMV(CAT)T7-SB100 (a gift from Zsuzsanna Izsvak, Addgene #34879),^57^ using FuGENE6 (Promega) in a 1:3 ratio with DMEM. On day 6, the cells were exposed to HEK media with 1 µg/mL doxycycline (Sigma) to induce expression of the BFP/iCasp9 from the landing pad. The cells were sorted with a BD FACSAria IIIU (BD Biosciences), using a 100 µm nozzle for cells with high levels of BFP and high levels of GFP (Figure S14B) into HEK media on amine-coated plates (Corning). Single-cell colonies were grown in HEK media, with the addition of 1 µg/mL doxycycline 48 hours before being screened on a BD LSRFortessa SORP (BD Biosciences) to identify colonies with high rates of both landing pad (*BFP*) and *KCNQ1* (*GFP*) expression (Figure S14C). Two distinct colonies with high blue and green fluorescence were expanded for further experiments. The landing pad cell line expressing WT KCNQ1 is referred to as LP-KCNQ1 and the cell line expressing KCNQ1 S140G is referred to as LP-KCNQ1-S140G.

For transient transfection experiments, plasmids that constitutively expressed *KCNE1* or *KCNQ1* were cloned using NotI restriction digestion into pIRES2:EGFP or pIRES2:dsRED2 to create all four combinations: p*KCNE1*:IRES:EGFP, p*KCNE1*:IRES:dsRED2, p*KCNQ1*:IRES:EGFP, and p*KCNQ1*:IRES:dsRED2 (see Supplemental Methods for more details).

### KCNE1 library creation

Each of the 129 *KCNE1* residues was comprehensively mutated using inverse PCR on the compact template plasmid.^23, 46^ For each mutagenesis reaction, the forward primers contained a 5’ degenerate NNK sequence for each codon to introduce all 20 sense amino acid variants and 1 nonsense variant at each site. The 129 PCR reactions were pooled, PCR-purified (QIAGEN), phosphorylated with T4 PNK (NEB), and self-ligated using T4 ligase (NEB). The pooled product was re-PCR-purified and electroporated into ElectroMAX DH10B Cells (ThermoFisher) using a Gene Pulser Electroporator (Bio-Rad; 2.0 kV, 25 µF, 200 Ω). Serial dilutions of the electroporated cells were plated on Ampicillin plates to determine transformation efficiency. The library was purified using a MaxiPrep kit (Qiagen) and subcloned into a promoterless AttB pIRES:mCherry-BlastR^18, 46^ plasmid using restriction digestion with AatII and Aflll (NEB). The library was then re-electroporated and purified as above.

For quantification, each variant construct was tagged with a random 18mer barcode to the library plasmid. The barcode was generated using ag289 (spacer-BsiWI-AflII-N18-AatII-XbaI-spacer) and ag290 (reverse complement of the 3’ end of ag289). The two primers were annealed by incubating at 95° for 3 minutes and cooling by 1°/10 s. The annealed primers were extended to fully double stranded DNA using Klenow polymerase (NEB). The barcode was purified by phenol-chloroform extraction, digested with BsiWI and XbaI (Thermo Fisher), and re-purified by phenol-chloroform extraction. The resulting DNA fragment was ligated into the *KCNE1*-HA plasmid library. The plasmid library was deep sequenced to link each barcode to its associated variant (see Supplemental Methods). A map of the final barcoded plasmid library (p*KCNE1*-HA library:pIRES:dsRed:BlastR) is provided in Figure S3A.

### Integration of library into landing pad cells

LP, LP-KCNQ1, or LP-KCNQ1-S140G cells were cultured in 10 cm dishes in HEK media until approximately 60% confluent. Cells were then transfected with 25 µg of p*KCNE1*-HA library:IRES:dsRed:BlastR and 5µg of site-specific recombinase, pCAG-NLS-HA-Bxb1 (a gift from Pawel Pelczar, Addgene #51271)^58^ using Lipofectamine 2000 (Thermo Fisher) in a 1:2 ratio.^18^ 6-8 hours after transfection, the cells were fed fresh media. The following day, cells were passaged using Accutase (Sigma) into two 10 cm dishes and fed media supplemented with 1 µg/mL doxycycline (Sigma) to induce expression of the landing pad. With successful integration of a single AttB-containing plasmid, the landing pad expresses only one *KCNE1*-HA variant, mCherry, and BlastR. 24 hours later, cells were singularized in place using 1 mL Accutase, quenched with 10 mL selection media (HEK media as above with 1 µg/mL doxycycline, 100 µg/mL blasticidin (Sigma), and 10nM AP1903 (MedChemExpress)). Addition of blasticidin and AP1903 selected for successfully integrated cells that expressed *KCNE1*-HA library:pIRES:mCherry:BlastR and against non-integrated cells that continued to express BFP/iCasp9 caspase. Cells were grown in selection media for 8 days post-transfection. For the LP-KCNQ1-S140G line, cells with successful integration were enriched in HEK selection media containing a high concentration (500 nM) of the HMR 1556 (Tocris), a specific I_Ks_ blocker with an IC of 10.5 nM.^59^

### Trafficking assay

We measured cell surface trafficking of the *KCNE1*-HA library in the presence and absence of *KCNQ1*, using LP-KCNQ1 and LP cells, respectively. Each experiment was performed in triplicate (three independently transfected, stained, sorted, and sequenced replicates).

To stain KCNE1-HA at the cell surface, day 8 post-selection, cells were dissociated using Accutase and quenched with HEK selection media. For any subsequent cellular suspension, cells were centrifuged for 5 minutes at 300 g and suspended in the new solution. The cells were washed with block solution (1% FBS +25 mM HEPES pH 7.0 [Sigma] in PBS without Ca^2+^/Mg^2+^). The cells were then stained with 1:500 anti-HA AF647 conjugated antibody (Cell Signaling #3444) in block solution for 45 minutes at room temperature in the dark and washed three times. Stained cells were resuspended in block solution at a concentration of 4-6 million cells/mL for sorting. Two trafficking experimental replicates were performed with cells in suspension as described above. One replicate was stained while cells were adhered on the dish (see Supplemental Methods).

Stained cells were FACS sorted on the BD FACSAria III (BD Biosciences) using a 100 µm nozzle. Compensation was performed using cells expressing single fluorophores or AF647 compensation beads (Thermo Fisher). Single cells were identified based on forward and side scatter signals. BFP^-^/mCherry^+^ single cells (with and without GFP for *KCNE1* with and without *KCNQ1* experiments, respectively) were divided into 4 groups of approximately equal size based on AF647 labeling (Figure S15). At least 800,000 cells per group were sorted and replated onto amine-coated plates containing HEK media. Cells were expanded and genomic DNA was extracted from each group with the Quick-DNA midiprep kit (Zymo). Q5 polymerase was used to amplify the barcodes from gDNA with 20 cycles of amplification using Illumina-compatible primer pairs (Table S2), and the barcodes were sequenced on the Illumina NovaSeq PE150 with at least 25 million reads per sample. Normalized cell surface trafficking scores were calculated from a weighted average of barcode counts from these samples (see Supplemental Methods for details).

### Cell fitness assay

The cell fitness assay was validated by 1:1 competition experiments on control variants (see Supplemental Methods for details) and then conducted on the entire library. The comprehensive library was integrated into the landing pad of cells as described above. Fitness of cells expressing *KCNE1*-HA variants in the library was measured in LP-KCNQ1-S140G in three independent replicates. 8 days after successful plasmid integration, termed Day 0 in subsequent analyses, cells were washed with selection media to remove HMR 1556, and a subset of cells harvested for gDNA extraction. Cells were passaged as needed. Additional samples were harvested for gDNA extraction on Day 8 and Day 20. Genomic DNA from these three time points (Days 0, 8 and 20) was isolated. Barcode pools were amplified and sequenced using the Illumina NovaSeq as described above.

### Population-level analyses

The data for putative controls were obtained from gnomAD v.2.1.1 (exomes and genomes, accessed October 2022).^54^ Variant annotations were obtained from the ClinVar database (accessed October 2022).^13^ In addition, we performed a literature review that consisted of a PubMed search (on November 30, 2022) for “KCNE1.” The resulting 929 manuscript titles and abstracts were manually reviewed to collate published patch clamp electrophysiology and patient data (Additional File 1, see Supplemental Methods for more details). Variants in at least 1 patient with LQT5 (Romano-Ward or one allele of JLNS) and an AF < 2.5×10^-5^ across all ancestries in gnomAD were classified as putative LQT5-associated variants. Similarly, the 2 variants previously seen in cases with atrial fibrillation, G25V and G60D, also had allele frequencies < 2.5×10^-5^, and thus were classified as putative atrial fibrillation-associated variants. Putative benign variants had an estimated penetrance of < 10% (#) and had an allele frequency of > 3×10^-5^ in gnomAD. Only 1 variant (G55S) with an allele frequency of > 3×10^-5^ had a penetrance of > 10% and was excluded.

### ACMG Assay Calibration

Missense variants classified in ClinVar as pathogenic/likely pathogenic or in our curated list of putative LQT5 variants were classified as “presumed cases.” Missense variants classified as benign/likely benign in ClinVar or in our curated list of putative normal variants were classified as “presumed controls”. Variants with score estimates (mean and 95% confidence interval) below the 2.5^th^ percentile or above the 97.5 percentile of the synonymous distribution were defined as loss-of-function and gain-of-function respectively. Variants with score estimate within the cutoffs were classified as “functionally WT-like.” Missense variants score estimate that overlapped the cutoffs were classified as “indeterminate.” Only one variant (Y107R) had a function score estimate higher than the 97.5 percentile cutoff. To determine the strength of evidence at which MAVE scores could be used as evidence for the ACMG functional assay criteria (PS3/BS3), the assay OddsPath was calculated using previously described methods.

## Additional files

Additional File 1: Literature review of KCNE1 variants. File name: AdditionalFile_LitRev.csv

Additional File 2: Function and trafficking scores for KCNE1 variants. File name: AdditionalFile_MAVEscores.csv

Additional File 3: Structure of the KCNE1/KCNQ1/calmodulin complex with mean function scores at each residue overlaid on KCNE1. File name: KCNE1-AF2-full-func-score-mapped.pdb

Additional File 4: Structure of the KCNE1/KCNQ1/calmodulin complex with mean trafficking scores at each residue overlaid on KCNE1. File name: KCNE1-AF2-full-traffick-score-mapped.pdb

Additional File 5: Computational scores curated for variants tested in this study. File name: AdditionalFile_CompScores.csv

