## Supplemental Information for "High-throughput functional mapping of variants in an arrhythmia gene, *KCNE1*, reveals novel biology"

### Supplemental Methods

***Modifying pIRES2 to have a NotI restriction site***

Using QuikChange mutagenesis, we modified the pIRES2-dsRED2 and pIRES2-EGFP mammalian expression constructs (Clontech) in two ways. We introduced a NotI restriction site in the multiple cloning site (MCS) and removed a NotI site outside the MCS. These plasmids were used for cloning *KCNE1* or *KCNQ1* cDNA into the MCS using NotI restriction digestion for expression studies.

***Subassembly of the variant library***

To subassemble the library, i.e., associate each 18-mer barcode with its specific *KCNE1* variant, the library was divided into 9 regions as shown in Figure S14D. The barcode was located 110 bp upstream of KCNE1 in the plasmid, so PCR primers were designed to amplify both the barcode and selected regions of the KCNE1 gene. Each region was sequenced on the Illumina NovaSeq platform using paired-end 150 base pair sequencing to give at least 100 million reads per region. The barcodes were extracted from reads using custom Python and Unix scripts. Each read included in the downstream analyses was required to have the correct anticipated 6 base prefix and 6 base suffix surrounding the barcode. Barcodes with over 500 total reads of subassembly data were retained, and barcodes were required to differ by at least 2 SNVs from each other. 99,418 candidate barcodes met these criteria (Figure S14E). The reads associated with each barcode were aligned to the reference *KCNE1*-HA plasmid sequence using the Burrows-Wheeler Alignment (bwa) tool and variants were called using samtools.^1,2^ Any barcodes associated with WT constructs, variants within the HA tag, multiple *KCNE1* variants, insertions or deletions were excluded from all analyses, resulting in 80,282 “good” barcodes each corresponding to a single nonsense, missense, or synonymous variant out of 2,592 unique protein variants.

***In-dish cell staining***

Cells in replicate 1 of the *KCNE1* with *KCNQ1* cell surface trafficking experiment were stained while adhered to the dish as follows. Cells were washed with OptiMEM (Thermo Fisher), before being stained in dishes for 30 min with 1:500 anti-HA AlexaFluor 647 antibody (Cell Signaling, #3444) dissolved in DMEM. The cells were washed with HEK media 3 times, and then briefly with OptiMEM. Next, the cells were incubated with 0.5 mM EDTA in PBS (Corning) for 7 minutes at room temperature and resuspended in DMEM. All other steps were identical to the "in-suspension" staining described in the Methods section. Since the trafficking scores from this replicate were highly concordant with those from the other two replicates where cells were stained in suspension (Figure S4B), the scores from all three replicates were combined and averaged as described below.

***Trafficking score calculation***

For each sample, custom Python, R, and Unix scripts were used to process reads and calculate trafficking scores as follows. Reads that matched the expected 6-base sequences before and after each barcode were retained and aggregated. Only barcodes in the list of 80,282 “good” barcodes from the subassembly (see above) were used for subsequent analyses.

The barcode counts were normalized to the total number of reads in each sample to calculate a "reads per million" metric according to the equation below:

$$F_{i,j}= \frac{R_{i,j}}{T_{j}} \times{10}^{6}$$

Where *i* is the barcode ID, *j* is the bin number (1 through 4, Figure S15A and S15B), *R_i,j_* is the number of raw reads for barcode *i* in bin *j*, and *T_j_* is the total number of reads in bin *j*.

Next, the read counts were aggregated by variant:

$F_{v,j}= \sum_{i=1}^{barcode} F_{i,j}$

Where *i* is the barcode ID corresponding to variant *v*, *j* is the bin number (1 through 4), *F_v,j_* is the frequency of the variant *v* in bin *j*, and *F_i,j_* is the frequency of barcode *i* in bin *j*.

A weighted average was then calculated for each variant (*W_v_*) as follows:

$W_{v}= \frac{{0\times F}_{v,bin 1}+ {1\times F}_{v, bin 2}+{2\times F}_{v, bin 3}+{3\times F}_{v, bin 4}}{F_{v,bin 1}+ F_{v, bin 2}+F_{v, bin 3}+F_{v, bin 4}}$

Where cells expressing variants with lowest and highest cell surface anti-HA labeling are in bins 1 and 4 respectively.

Next, the weighted averages were linearly transformed so that synonymous variants would have a median score of 1, and trafficking-null nonsense variants would have a median score of 0. This way, the normalized scores could be biologically interpreted: for example, a variant with a score of 0.5 would traffic approximately half as well to the cell surface compared to WT. Since we observed that the nonsense mutations followed a bimodal distribution, with a drastic shift in scores at residue #56, this normalization of raw scores was based on the distribution of synonymous and early (i.e., before residue 55) nonsense variants, as follows:

$$S_{v}= \frac{W_{v}-Median(W_{v,syn})}{Median\left( W_{v,syn} \right)-Median(W_{v,early nonsense})}$$

Where for variant *v*, *S_v_* is the normalized score, *W_v_* is the weighted average across the 4 bins, and *W_v,syn_* is the set of weighted averages of all synonymous variants represented in the library. *S_v_* for each variant was calculated across each replicate. For quality control (QC), the frequency of each variant across each replicate experiment was calculated as follows:

$$F_{v,tot}= F_{v,bin 1}+ F_{v, bin 2}+F_{v, bin 3}+F_{v, bin 4}$$

For different cutoffs of *F_v,tot_*, the coefficient of variance (CV) for the distribution of synonymous variants was calculated across each replicate as follows:

$$CV=\frac{std dev(S_{v,syn})}{mean(S_{v,syn})}$$

For each replicate, we plotted the CVs, the mean scores of the synonymous distribution, and the number of unique missense variants against different cutoffs of *F_v,tot_* (Figure S4A). The threshold for *F_v,tot_* to minimize the CV and maximize the number of unique missense variants was determined to be 100 for each replicate for downstream analyses. Accordingly, variants with a total frequency lower than 100 in each replicate were dropped from trafficking score calculations. 39/2,592 variants did not meet this frequency threshold. For each variant meeting this QC threshold, the normalized scores *S_v_* were averaged across 3 replicates and standard error of the mean calculated. Cutoffs for trafficking increase and loss were defined as mean ± 1.96xSD of the synonymous variant scores, i.e., 97.5^th^ and 2.5^th^ percentile respectively. Variants with score estimates (mean and 95% confidence interval) below the 2.5^th^ percentile and above the 97.5^th^ percentile of the synonymous distribution were defined as loss-of-trafficking and gain-of-trafficking respectively.

***Fitness Score Calculation***

To calculate fitness scores, custom Python, R, and Unix bash scripts were used to process reads for each replicate, where each replicate was harvested on Days 0, 8 and 20. Reads were filtered to remove mismatches in the expected 6-base region before and after the barcode position and aggregated by barcode. Only barcodes found in the list of 80,282 “good” barcodes from the subassembly were used for downstream analyses as described above.

Similar to the trafficking score calculation, barcode counts were normalized to the total number of reads in each sample to calculate a "reads per million" metric according to the equation below:

$$F_{i,d}= \frac{R_{i,d}}{T_{d}} \times{10}^{6}$$

Where *i* is the barcode ID, *d* is the day number (0, 8 or 20), *R_i,d_* is the number of raw reads for barcode *i* in the sample harvested on day *d*, and *T_d_* is the total number of reads in the sample harvested on day *d*.

Next, the read counts were aggregated by variant:

$F_{v,d}= \sum_{i=1}^{barcode} F_{i,d}$

Where *i* is the barcode ID corresponding to variant *v*, *d* is the day number (0, 8 or 20), *F_v,d_* is the frequency of the variant *v* in the sample harvested on day *d*, and *F_i,d_* is the frequency of barcode *i* in the sample harvested on day *d*.

Next, a log_2_ transformed ratio of the frequency of each variant on day 0 compared to day 20 was calculated (*R_v,20_*) as below:

$$R_{v,20}={log}_{2}\left( \frac{F_{v,20}}{F_{v,0}} \right)$$

A similar ratio (*R_v,8_*) representing the function scores on day 8 was also calculated. Next, day 8 and day 20 scores were linearly transformed as above. Synonymous variants would have a median score of 1, and trafficking-null early nonsense variants would have a median score of 0. 54/2,592 variants with less than 30 reads at day 0 of the fitness assay (*F_v,0_*) were dropped from score calculations.

Heatmaps were generated using the R package heatmap.2^3^ and ROC curves were generated using plotROC.^4^

***Cell fitness assay — control variants***

To determine the degree of selection against control variants, a 1:1 ratio of single *KCNE1* variant and non-*KCNE1* containing AttB plasmids was integrated into the landing pad of LP-KCNQ1-S140G cells as described in the Methods and Figure 3B. During initial plasmid integration and selection for successful integration events, cells were grown in 500 nM HMR 1556 (Tocris) to inhibit I_Ks_. The KCNE1^-^ plasmid consistently results in a lower red fluorescence level than the KCNE1^+^ plasmids and allows for distinguishing between cell populations. After 8 days to enrich for successfully integrated cells, HMR 1556 was removed from cell media to allow K^+^ flux-based selection (Day 0). Cells were grown and passaged as described above. The ratio of KCNE1^-^ (low dsRed) to KCNE1 variant (high dsRed) cells was serially measured by flow cytometry (LSR Fortessa SORP, BD Biosciences). These ratios were then normalized to the ratio at Day 0 corresponding to HMR 1556 removal. This experiment was performed with 3 replicate samples per variant. The mean and standard error of the three replicate measurements was calculated for each data point.

***Cell staining for manual flow cytometry validation and microscopy***

Several previously studied variants were examined by flow cytometry and confocal microscopy to validate the trafficking assay (Figures 1D and 1E). HEK293T cells were transfected with p*KCNQ1*:IRES2:dsRED2 and p*KCNE1*-HA:IRES2:dsRED2 using Lipofectamine 2000. 48 hours later, transfected cells were harvested using Accutase, washed in block (1% FBS +25 mM HEPES (Sigma, pH 7.0) in PBS without Ca^2+^/Mg^2+^), and incubated for 30 minutes with the anti-HA AlexaFluor 647 (AF647) antibody. If used for flow cytometry, the cells were washed three times and analyzed on the LSRFortessa SORP (BD Biosciences). If used for microscopy, the cells were washed, fixed in 4% paraformaldehyde (Thermo Fisher), and permeabilized in 0.2% SDS (Sigma, #L4390) for 5 minutes. Cells were stained with anti-HA AlexaFluor 488 (AF488) antibody (Cell signaling, #2350), washed and stained with 1:1000 Hoechst, and resuspended in 50% glycerol before being added to cover slips. Images were acquired on an inverted confocal microscope (Olympus) equipped with a spinning disk (Yokogawa), 60x silicone objective, and ORCA-Fusion CMOS camera (Hamamatsu). Excitation was achieved using 100 mW solid-state diode lasers. Hoechst was excited at 405 nm, AF488 at 488 nm, dsRED2 at 561 nm, and AF647 at 640 nm. 16-Bit images were acquired sequentially and exported to ImageJ (FIJI) for analysis. Images are shown with equivalent intensity scaling after background subtraction.

***Literature Review (additional information)***

Variants with an AF >0.1% in gnomAD were excluded from the literature review. For variants reported in multiple papers from the same site or with overlapping authors, we included patient data only from manuscripts with the largest number of reported patients to prevent double counting. Patient phenotype annotations were as reported in the manuscripts (i.e., we did not attempt to re-adjudicate the phenotype if a patient was annotated as having long QT syndrome). If only a QT interval without assertion of either long QT syndrome or unaffected carrier was given, we used a QTc of ≥460 ms to determine LQT5, as per a large *KCNE1* patient curation study.^5^ For patch clamp data, the peak current densities of *KCNE1* variants coexpressed with *KCNQ1* (normalized to WT) were collected. When possible, the current density at +60 mV was used, however other voltages (e.g., +40 mV) were used if those were the only data reported. In addition, we curated voltage of half activation (V_½_) and the time constant (τ) of deactivation (each normalized to WT). Some papers reported *KCNE1* variants that, when co-expressed with *KCNQ1*, had KCNQ1 only-like currents (i.e., complete loss of KCNE1 function) but did not quantify peak current. These variants were annotated as having a normalized peak current of 10% (the typical approximate peak current of KCNQ1 only compared to KCNQ1+KCNE1). For variants with multiple papers reporting *in vitro* patch clamp data for a variant, these data were averaged. For most literature reports, there is no record of whether the WT allele was S38 or G38. However, S38G does not substantially alter properties of the I_Ks_ channel^6^ and does not affect the QT interval in large, well-powered GWAS studies,^7^ and is therefore unlikely to significantly affect mutation results.

***Modeling KCNE1 structure with AlphaFold2***

We leveraged multiple deep-learning protein structure prediction algorithms (ESMFold,^8^ trRosetta,^9^ and AlphaFold2^10^) to generate tertiary structure models of KCNE1. The AlphaFold KCNE1 structural model was constructed through the AlphaFold Google Colab notebook, a slightly simplified version of the full AlphaFold2 algorithm.^11^ We ran the Colab notebook with default parameters using the full-length KCNE1 amino acid sequence only (UniProtKB: P51382) as well as both the full-length sequences of KCNQ1 (UniProtKB: P51787 isoform 1) and KCNE1, i.e., AlphaFold-Multimer.^12^ The latter was to test the hypothesis that KCNQ1 might influence KCNE1 conformation upon binding of the two proteins. It has been noted that the Colab notebook has a small drop in average accuracy for multimers compared to local AlphaFold installation.^13^ We compared the structures to each other and to a previously resolved NMR structure (PDB ID: 2K21; Figure S8).^14^

Comparing the resolved cryo-EM structure of KCNQ1/KCNE3 to the predicted models of KCNE1, the AlphaFold-Multimer model appeared to be the most biologically meaningful, and thus all further analyses used this model. We modeled the structure of KCNE1 in complex with KCNQ1 and calmodulin via protein-protein docking using the Rosetta macromolecular modeling software suite.^15,16^ Briefly, we first energy-minimized AlphaFold2 predicted KCNE1 structure with respect to the RosettaMembrane energy function.^17^ We then prepared the starting pose of KCNE1 with respect to tetrameric KCNQ1 by aligning energy-minimized KCNE1 structure to KCNE3 in the KCNQ1-KCNE3-calmodulin cryo-EM structure (PDB ID: 6V00) and removing the KCNE3 subunits after aligning. AlphaFold2 predicts a “low” to “very low” confidence flexible linker between the extracellular alpha helix and the transmembrane domain. This flexible linker causes KCNE1 residues 1-30 to clash with the transmembrane domains of KCNQ1 in the homology model. Thus, we removed residues 1-30 from the N terminus of KCNE1. In protein-protein docking, we treated KCNE1 as the moving component and set the values of perturbation flags as -dock_pert 0.3 0.8, -dock_mcm_trans_magnitude 0.1, and -dock_mcm_rot_magnitude 1.0 to enable more local sampling. We generated 5000 models of KCNE1 in complex with KCNQ1 and calmodulin with good convergence (Figure S16) and selected the model with the best interface score for structural analysis in this work. Models of the I_Ks_ complex with mean function and trafficking scores overlaid are given in Additional Files 3 and 4 respectively.

***Computational and evolutionary metrics***

We tested the performance of MAVE scores against previously published and validated computational metrics (Table S4).^18–33^ These consisted of metrics that assess variant deleteriousness on protein function and/or evolutionary conservation of the residues. These scores for each variant in KCNE1 were either obtained from dbNSFP version 4.0a,^34,35^ a database of functional prediction and annotation of all nonsynonymous SNVs, or calculated as previously described. A curated list of computational scores is provided in Additional File S5.

### Supplemental Figures

#### Figure S1: Overview of multiplexed assays of variant effect in this study

Comprehensive KCNE1-HA variant libraries were integrated into either LP-KCNQ1 cells (trafficking assay), LP cells (trafficking assay in the absence of KCNQ1) or LP-KCNQ1-S140G cells (fitness assay). Cells for the trafficking assays were stained with anti-HA antibody and flow sorted into four groups based on KCNE1 cell surface expression. Each group was deep sequenced to quantify variant presence in each group for trafficking score calculations (See Supplemental methods). Cells for the fitness assay were grown for 20 days and samples at Day 0, Day 8 and Day 20 were deep sequenced to calculate function scores.

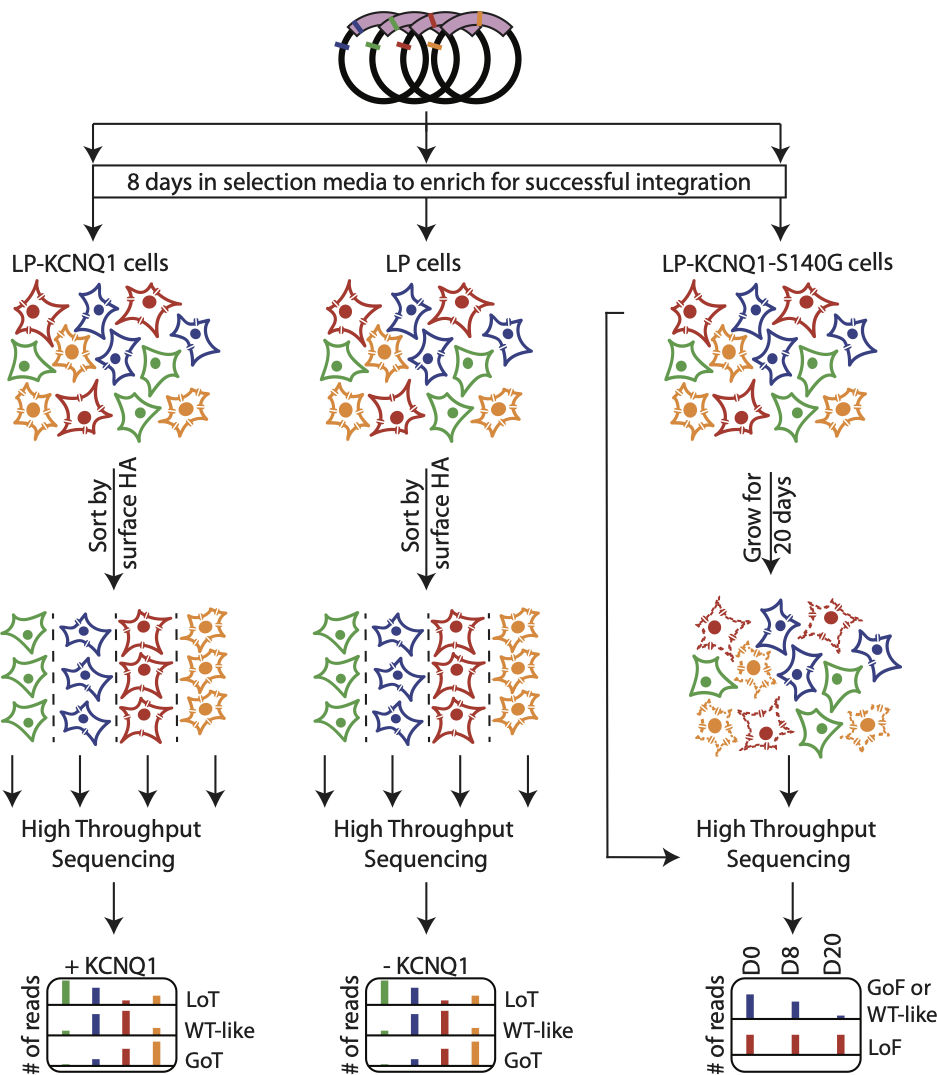

#### Figure S2: Quantification of KCNE1-HA protein at the cell surface by flow cytometry

A) Comparison of anti-HA labeling of two KCNE1 HA tags in alternate positions coexpressed with KCNQ1. An HA tag cloned between KCNE1 residues 34 and 35 (pink, this study) labeled approximately 3-fold more strongly with an anti-HA AlexaFluor 647-conjugated antibody than a previously-studied HA tag between residues 22 and 23 located in a predicted alpha helix (purple).^36^ B) Cell surface abundance of KCNE1-HA (34-35) for WT and previously studied variants. Data are from the same experiment in Figure 1E. Three independent replicates for each sample were transfected and stained. Blue = KCNE1-HA variants integrated into LP cells; Red = KCNE1-HA variants integrated into LP-KCNQ1 cells. Cells were stained with an Alexa Fluor 647-conjugated anti-HA antibody. For each sample, the median AlexaFluor 647 value across all cells is plotted. Bars show the mean of the 3 replicate values ± standard error. All values are normalized to the mean values for empty vector controls (0%) and KCNE1-HA-WT + KCNQ1 (100%). Co-expression of KCNQ1 results in a 6.1-fold increase in KCNE1-HA-WT cell surface abundance, but variant effects were similar in the presence and absence of KCNQ1.

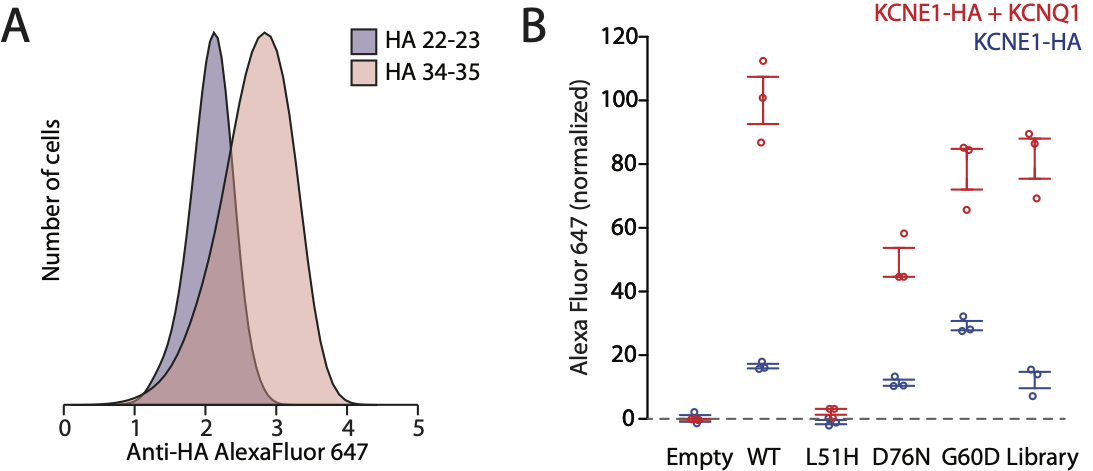

#### Figure S3: Creation, diversity, and coverage of the KCNE1 variant library

A) Creation of the KCNE1 variant library. Saturation PCR mutagenesis using degenerate NNK primers on a small, promoterless plasmid template was used to generate the variant library. The library was then subcloned with restriction digestion into the AttB landing pad compatible plasmid backbone. A poly-N 18 base barcode was added to the AttB plasmid by restriction digestion. B) Number of variants present subassembled per KCNE1-HA residue (max = 21: 19 missense, 1 synonymous, and 1 nonsense). The HA tag (not targeted during mutagenesis) had 2 off-target variant barcodes; these barcodes were discarded. C) Mean number of barcodes per variant at each KCNE1 residue. The ribbons represent the standard deviation. The HA tag is not represented in this panel.

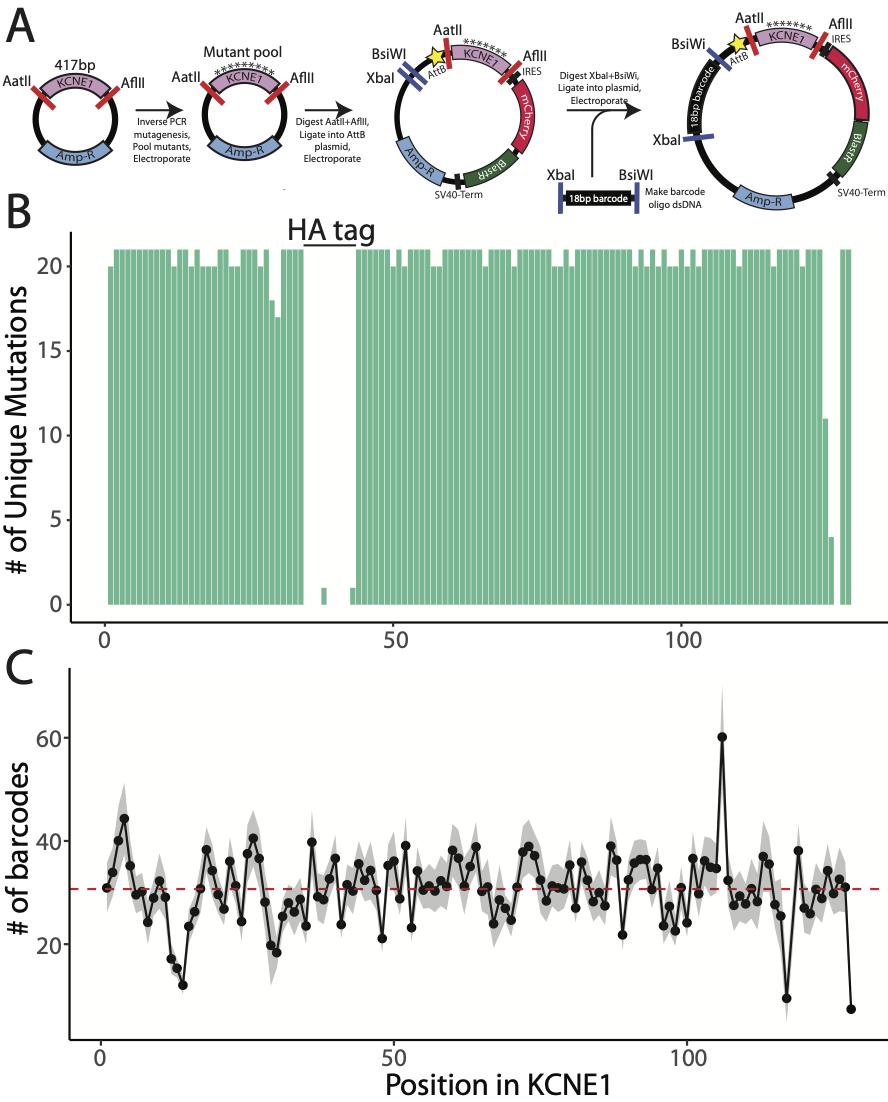

#### Figure S4: Inter-replicate distributions of KCNE1 trafficking scores (-KCNQ1)

A) Quality control prior to trafficking score calculation across three replicates. The coefficient of variation of the synonymous distribution (top), and the mean and median (middle, black and red respectively) synonymous variant scores as a function of total variant frequency, *F_v,tot_*, across each replicate experiment. The number of unique missense variants that passed each variant frequency cutoff (bottom). A cutoff of 100 (dotted line) was determined to minimize the coefficient of variation and maximize the number of unique missense variants for which scores were calculated. Variants with less than 100 reads per million in any replicate were excluded. B) inter-replicate Spearman correlation of trafficking scores based on variant category. ***: p-value < 0.001; **: p-value < 0.01; *: p-value < 0.05. For the trafficking scores, early and late nonsense variants are defined as variants at residue 1-55 and 56-129 respectively. C) Trafficking score correlations against reported measurements of variant cell surface trafficking. See Additional File 1 for the literature review dataset.

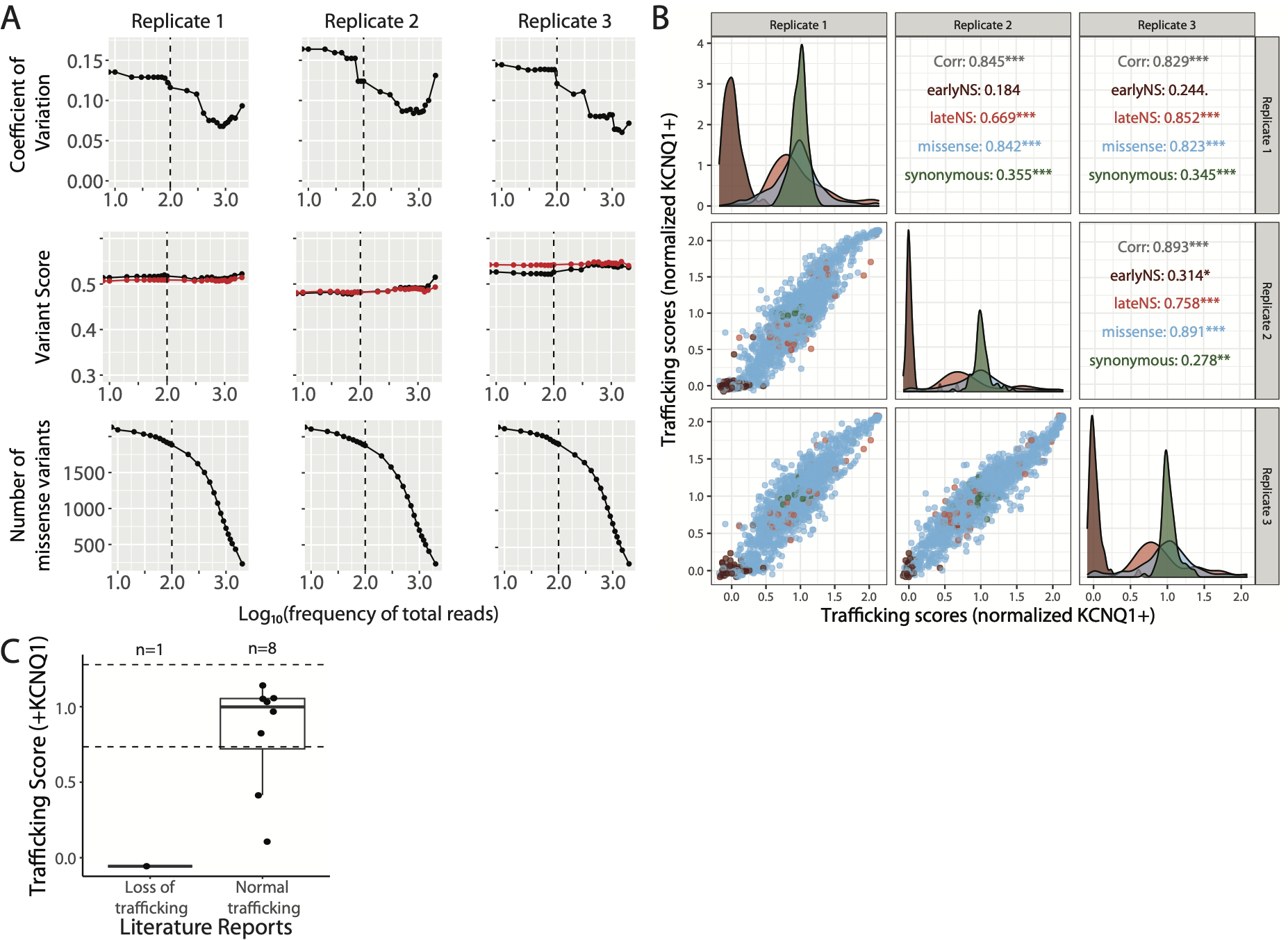

#### Figure S5: Cell surface expression heatmap for KCNE1 library (+KCNQ1)

A) Multiplexed assay of KCNE1 variant trafficking as seen in Figure S1. A comprehensively mutated barcoded plasmid pool was integrated into LP-KCNQ1 ("+KCNQ1"; panels B-C) or LP cells ("-KCNQ1", panels D-F). Cells were stained and sorted by surface HA and deep sequenced. B,E) KCNE1 trafficking heatmaps. Red, white, and blue indicate loss-of-trafficking, normal trafficking, and gain-of-trafficking, respectively. WT amino acids are indicated at each position with a dot. The colored ribbon indicates secondary structure as seen in Figures 2 and 4. C,D) For each residue, the proportion Gain-of-Trafficking (GoT) and Loss-of-Trafficking (LoT) missense variants is displayed. F) The number of unique variants observed in the -KCNQ1 trafficking map per position (max = 21).

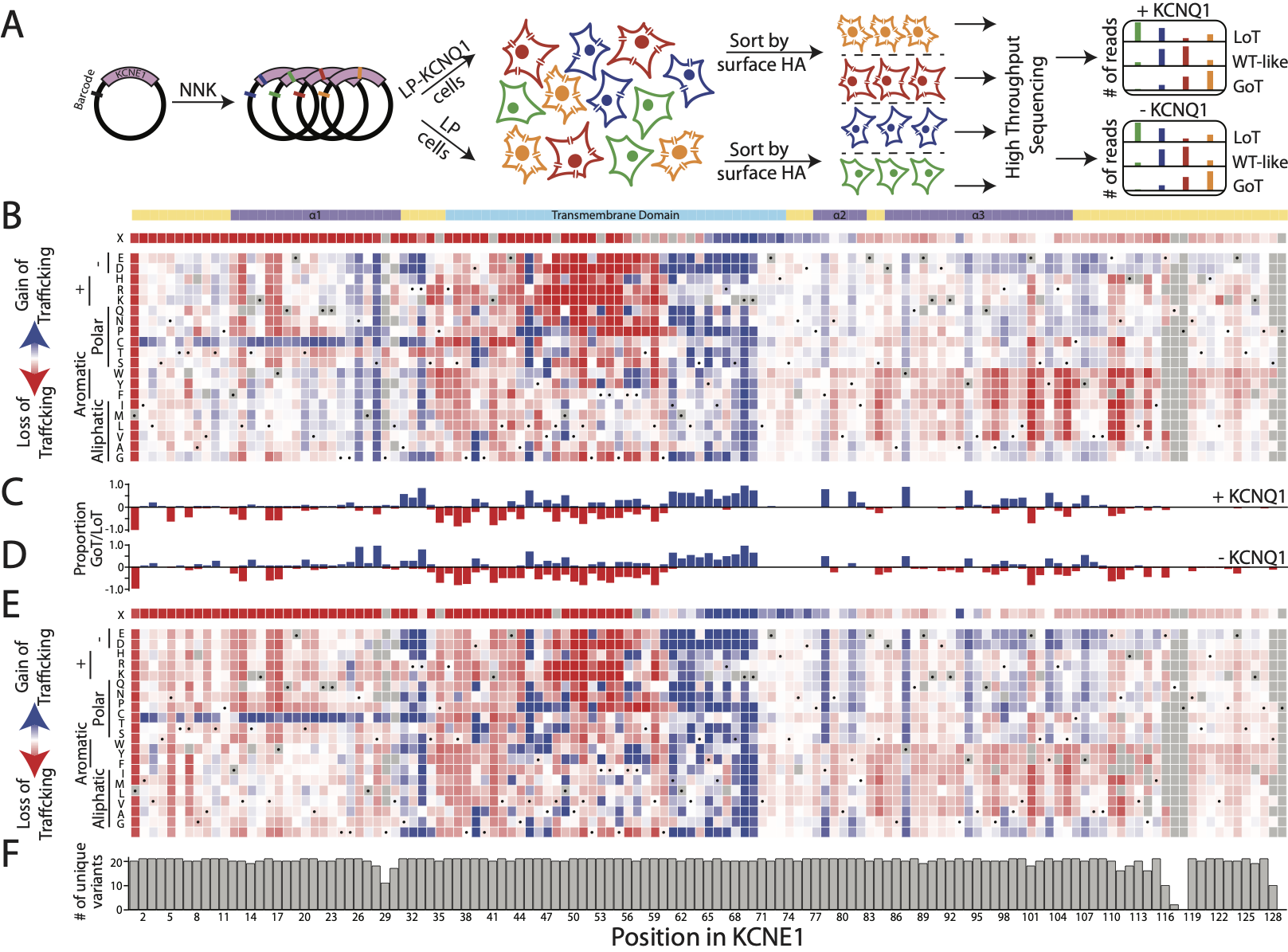

#### Figure S6: Development of function assay and distribution of function score

A) Variant abundances in the KCNE1 functional assay at various timepoints (in the subassembly/original library, at day 0, day 8, and day 20 of the growth experiment). Variant frequencies are expressed as normalized reads per million. B) Proportion of sequencing reads per variant category across different sequencing experiments: original subassembly, trafficking assay (conducted in KCNQ1-WT; sequenced at day 8), fitness assay (conducted in KCNQ1-S140G; sequenced at day 8 and day 20). C) Quality control prior to function score calculation across three replicates. The coefficient of variation of the synonymous distribution (top), and the mean and median (middle, black and red respectively) synonymous variant scores as a function of variant frequency at day 0, across each replicate experiment. The number of unique missense variants that passed each variant frequency cutoff (bottom). A cutoff of 31.6 (10^1.5^, dotted line) was determined to minimize the coefficient of variation and maximize the number of unique missense variants for which scores were calculated. Variants with less than 31.6 reads per million at day 0 in any replicate were excluded. D) inter-replicate Spearman correlation of functional scores based on variant category. ***: p-value < 0.001; **: p-value < 0.01; *: p-value < 0.05. Early and late nonsense variants are defined as variants at residues 1-55 and 56-129 respectively.

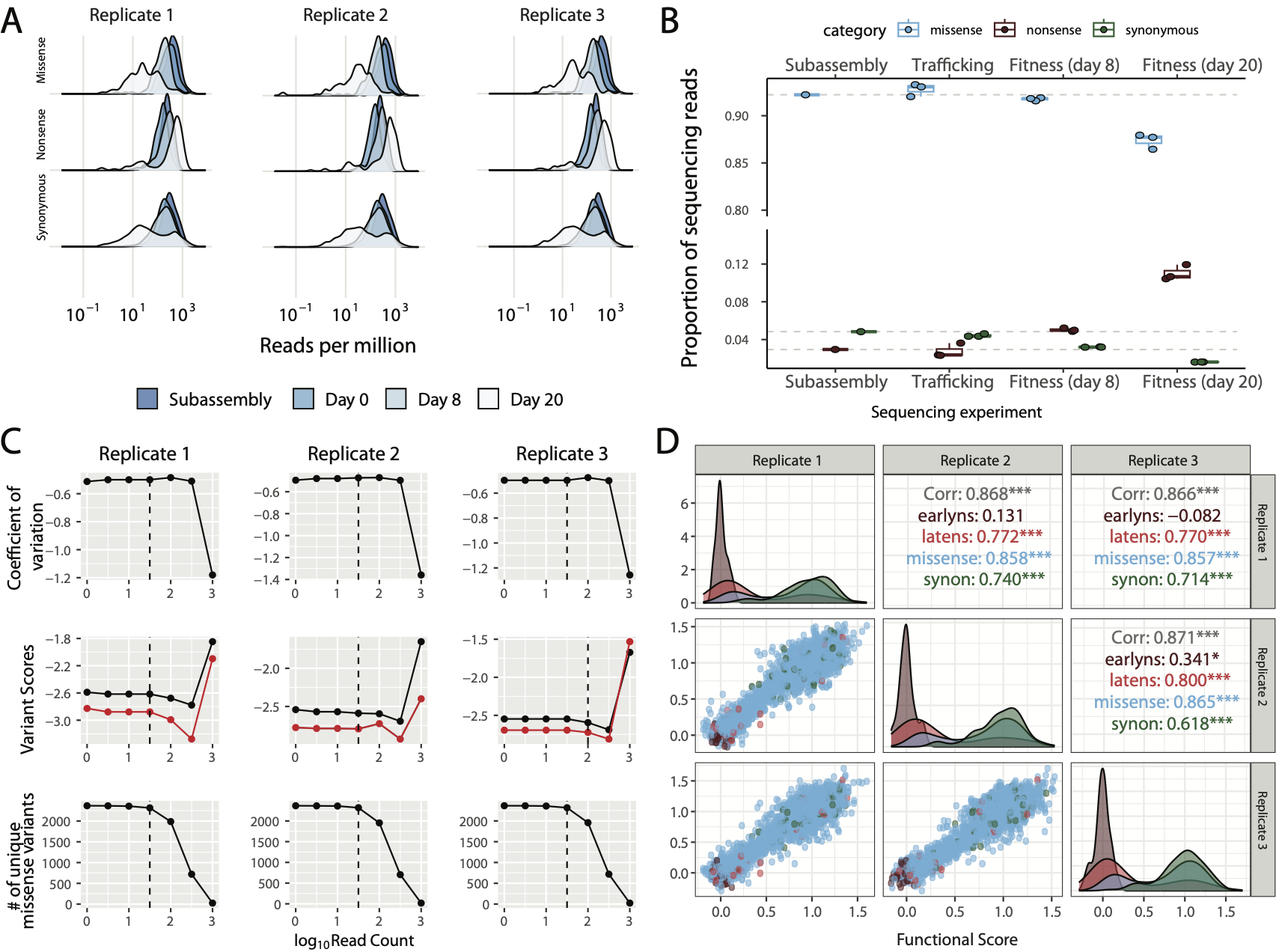

#### Figure S7: Relationship of function and trafficking variant scores and validation of gain-of-function variants

A) Distribution of nonsense variant function and trafficking scores show three categories of nonsense variants: those with trafficking and functional defects (brown, residue 1-55), those with WT-like trafficking but gating defects (red, residue 56-104) and those with WT-like trafficking and function (pink, residue 105+). B-C) Variants at glycosylation sequons alter trafficking in the absence of KCNQ1 but do not affect function scores. D) Peak and tail potassium currents of WT KCNE1-HA and two variants with the highest functional scores. Y107R was the only variant classified as gain-of-function classification. C106L, the variant with the second-highest function score, was classified as "indeterminate" since its score estimate overlapped with the 97.5^th^ percentile cutoff of function scores.

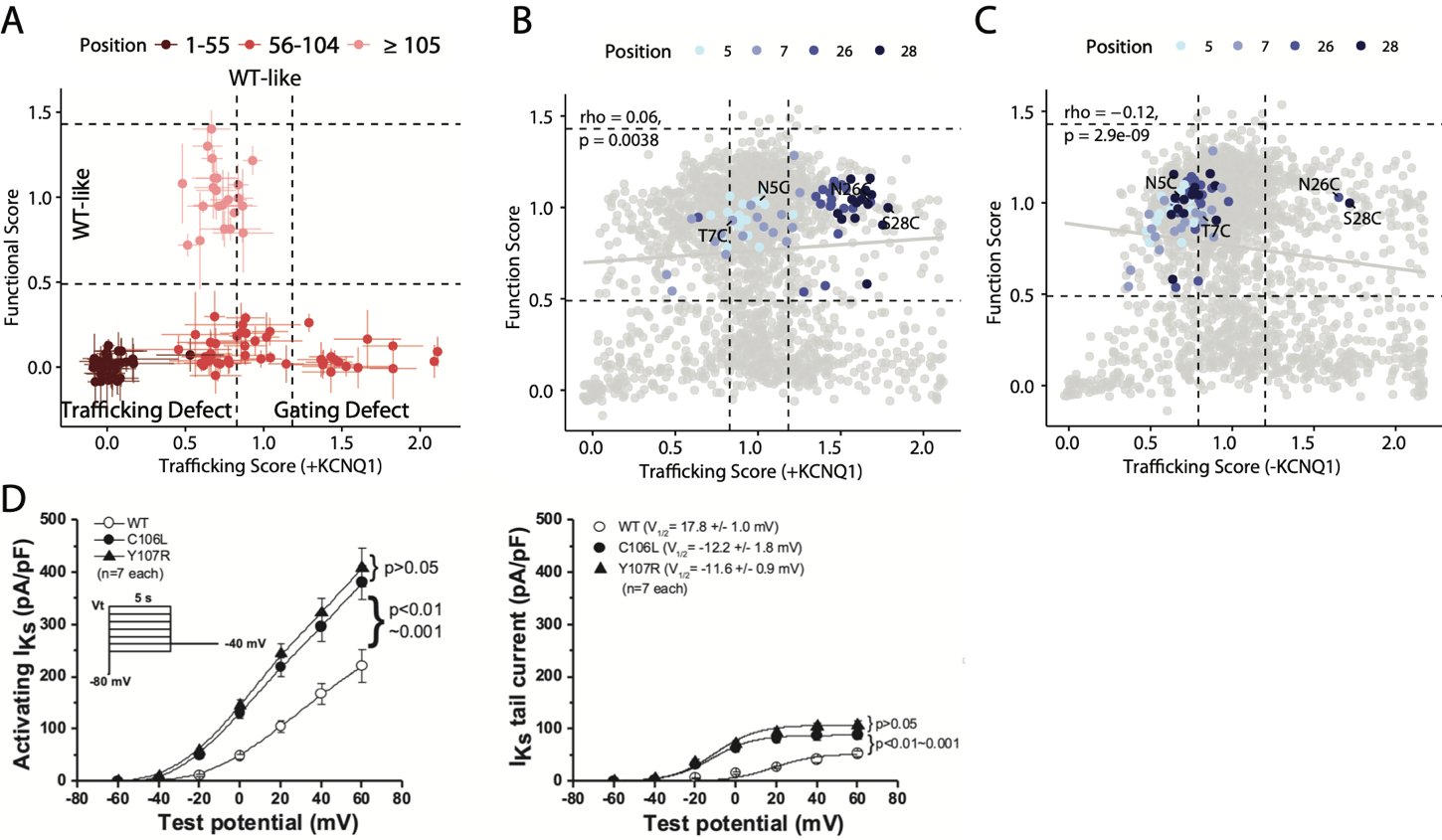

#### Figure S8: Comparison of structural models of full-length KCNE1 from various structure prediction methods and NMR

The image of each model generated by ESMFold,^8^ trRosetta,^9^ and NMR (PDB ID: 2K21)^14^ was rendered after alignment with the model predicted using AlphaFold-Multimer. After considering these structures in light of the KCNQ1:KCNE3 homology model, the AlphaFold-Multimer structure was selected as the primary structure for structural analyses in this study.

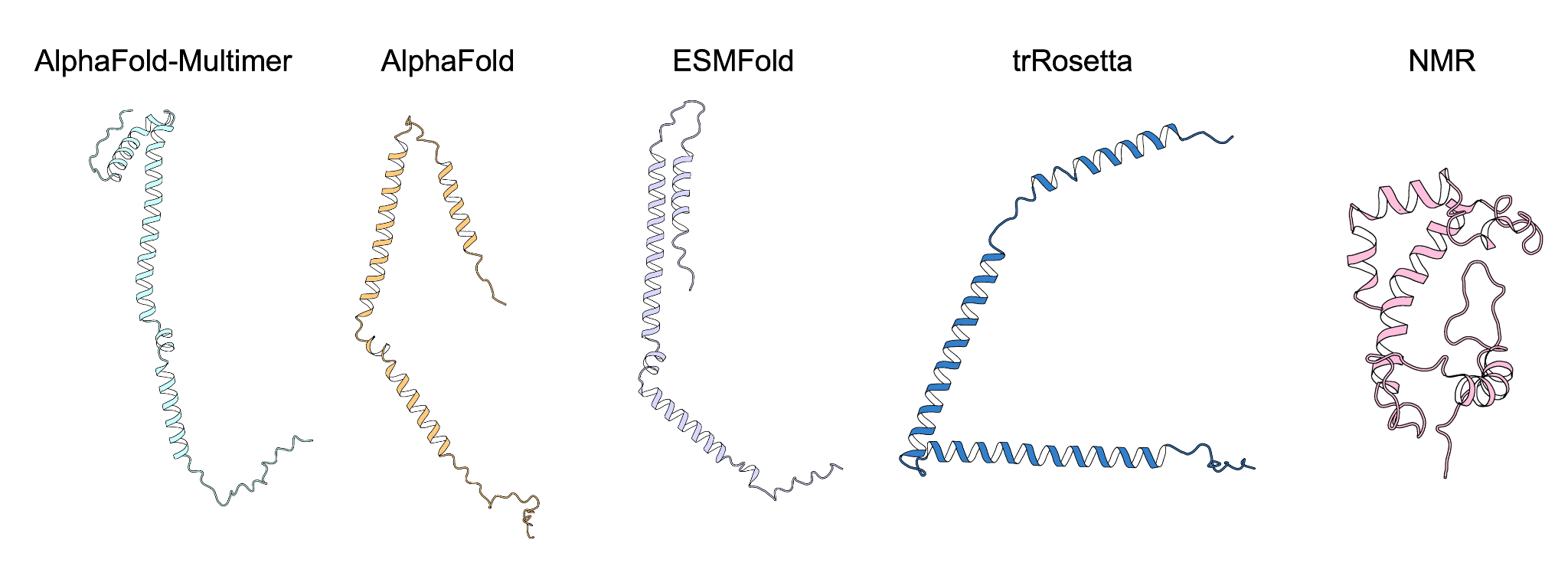

#### Figure S9: Correlation of function and trafficking scores with electrophysiological parameters and genomic variant categories

A-B) In addition to their strong correlation with peak current (Figure 6A), function scores also correlate with other electrophysiological properties of the I_Ks_ channel (A: V_½_ activation deviated from WT, B: deactivation time constant, as a percent of WT). C-E) Trafficking scores are less correlated with electrophysiological properties (C: peak current, D: V_½_ activation, and E: deactivation time constant) than function scores . F) Variants achievable by a single SNV (blue) are less likely to be trafficking-deficient than all other variants (black; p = 7.4x10^-5^, Wilcoxon test). G-H) Variants achieved by genomic transitions (black) are less likely to be functionally deleterious (G) than variants achieved by genomic transversions (gray; p = 2x10^-4^, Wilcoxon test). The effect does not exist in trafficking variants (H; p = 0.973, Wilcoxon test).

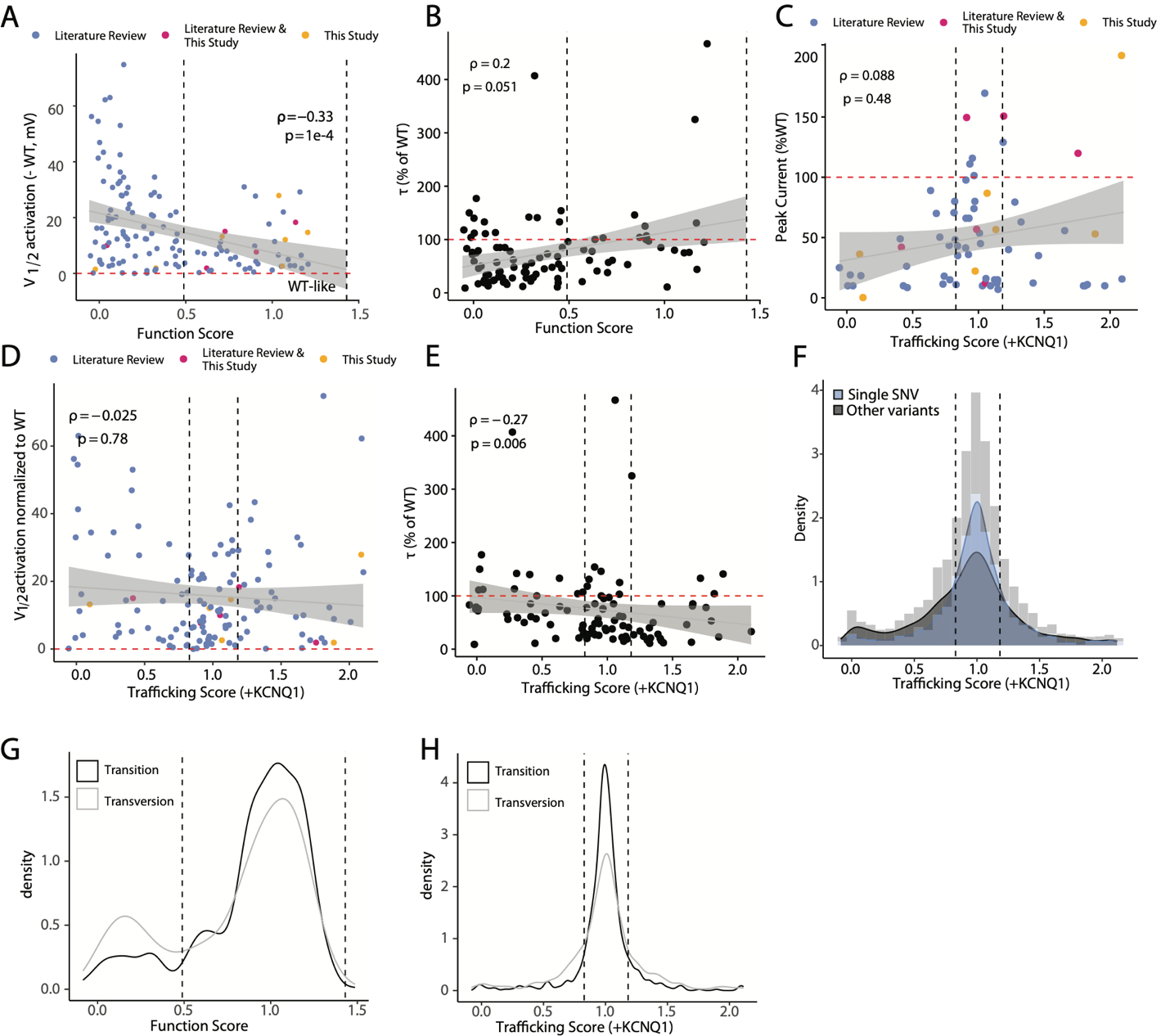

#### Figure S10: Distribution of function scores by gnomAD category and MAVE data prediction performance

A) Variants more common in gnomAD are more likely to have WT-like function scores. B) Receiver operator characteristic curves evaluating prediction of variant pathogenicity based on computational metrics and MAVE data. MAVE function scores perform equally as well as previously developed computational metrics of protein function and evolution.

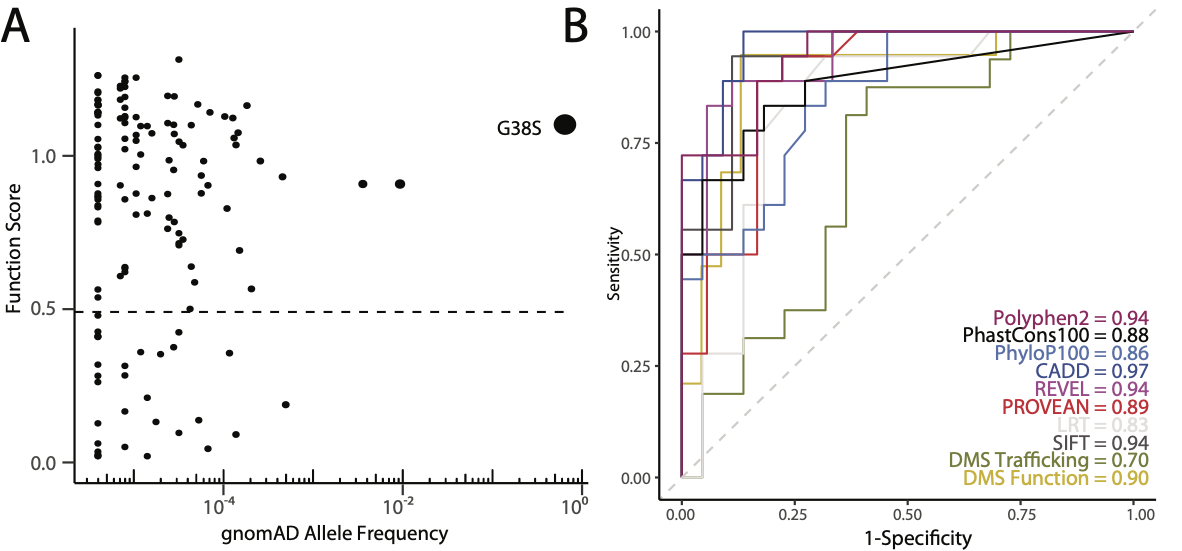

#### Figure S11: Spearman correlation of function scores to previously developed computational metrics of protein function and evolution

Correlation coefficients and p values are given. Blue dotted line represents the trend line.

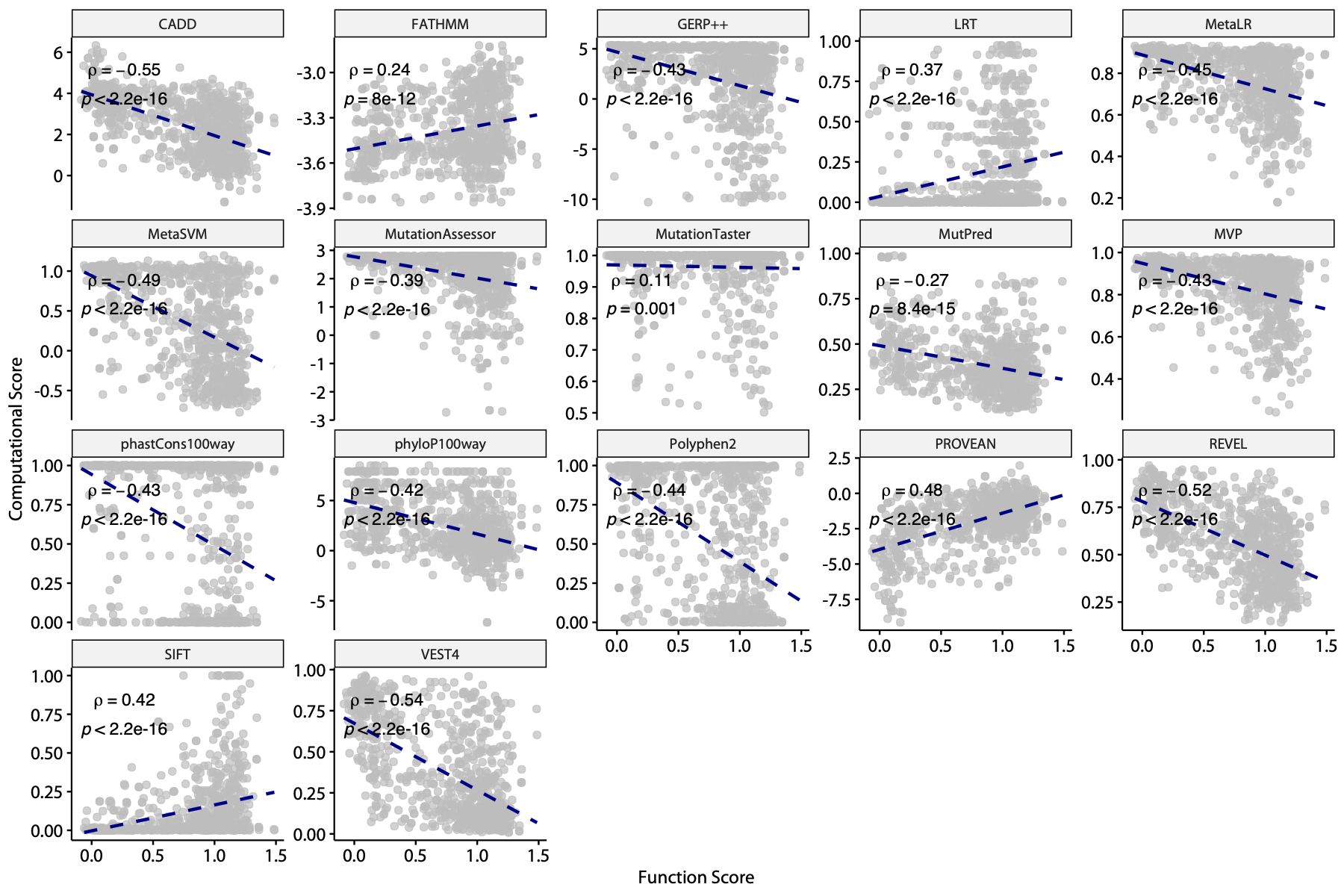

#### Figure S12: Correlation of trafficking scores to population and clinical case/control data

A) Variants more common in gnomAD are more likely to have WT-like trafficking scores. B) Distribution of trafficking scores of variants present in and absent from gnomAD (black and grey, respectively). Variants absent from gnomAD are slightly more likely to have deleterious trafficking scores (p = 4.2x10^-3^, Wilcoxon test) C) Trafficking score distribution of variants annotated in ClinVar by category. 3/4 P/LP variants have deleterious scores, and 5/7 B/LB variants have WT-like trafficking scores. D) Trafficking score distributions of presumed control and disease-associated variants.

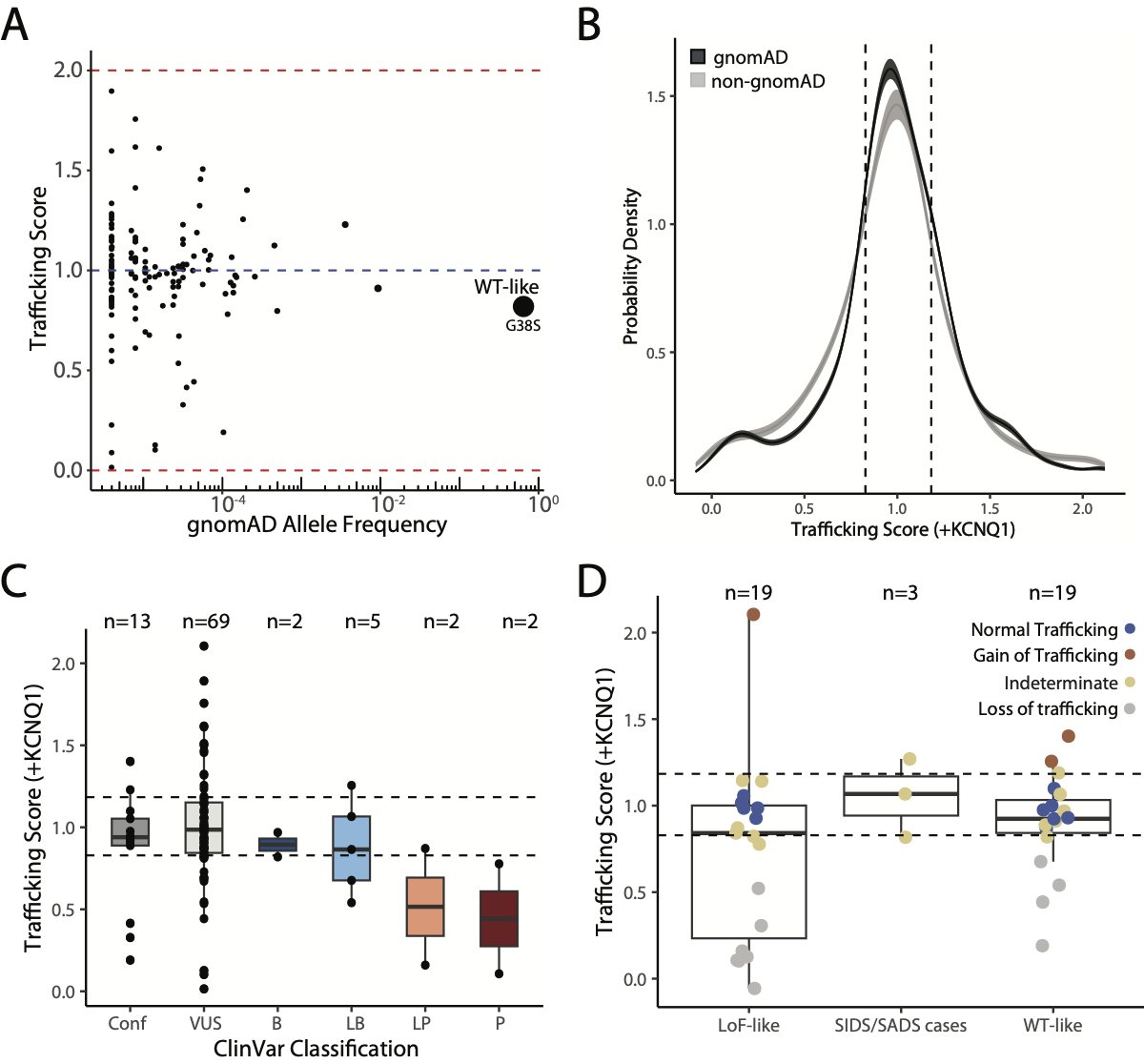

#### Figure S13: Spearman correlation of trafficking scores to previously developed computational metrics of protein function and evolution

Since gain-of-trafficking variants have been associated with a deleterious phenotype, i.e., atrial fibrillation, we checked the correlation of computational metrics to deviation of trafficking score from WT. Spearman correlations and p values are given. Blue dotted line represents the trend line.

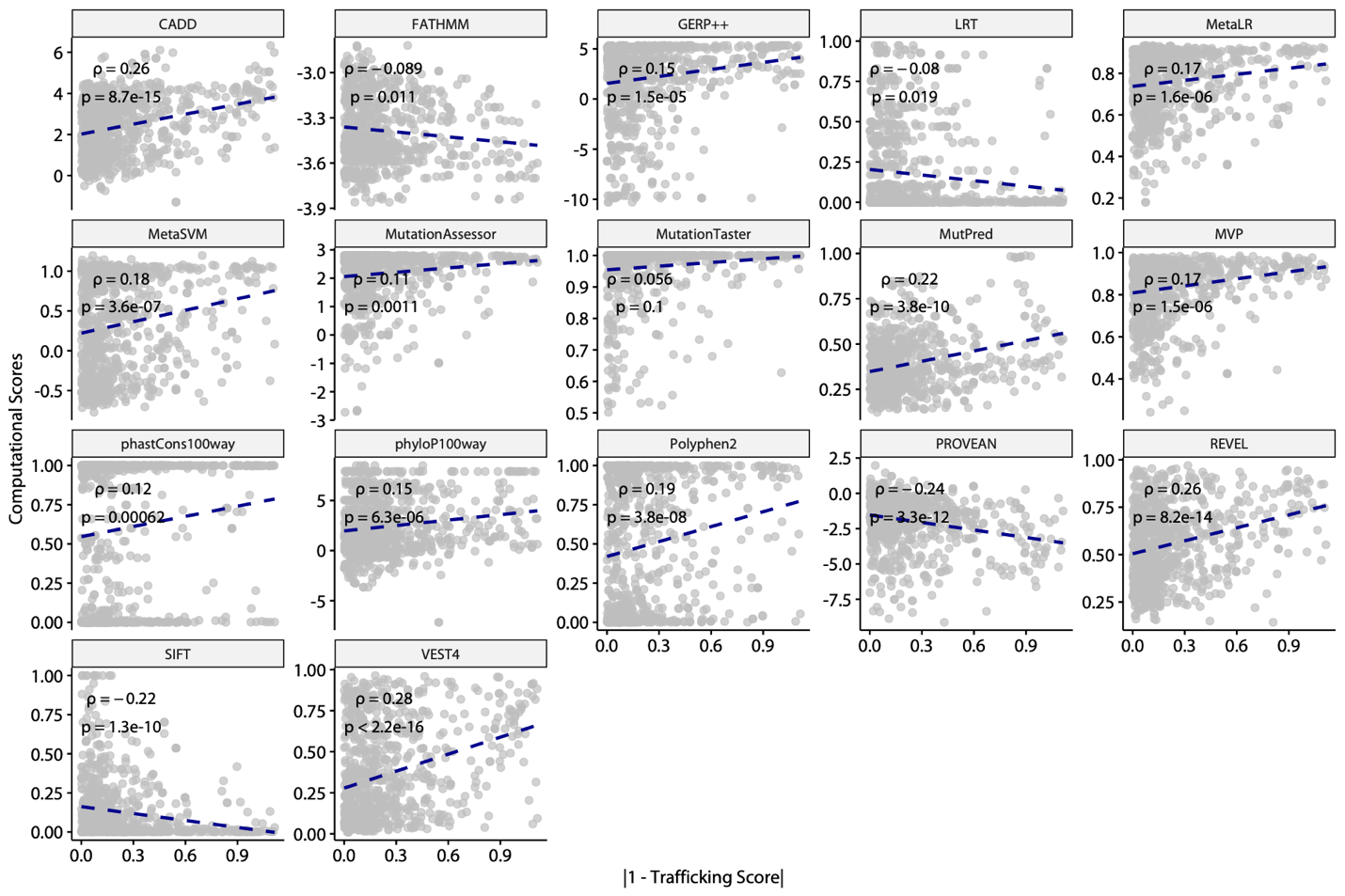

#### Figure S14: Plasmid maps, flow cytometry gates, and sequencing procedures in the study

A) Three different plasmids, constitutively expressing *KCNE1* (left), landing pad-compatible AttB-*KCNE1*-mCherry-BlastR (middle), and non-expressing minimalist *KCNE1* (right) were used for cloning in the study. B-C) Representative fluorescent marker expression and gating to select for LP-KCNQ1 cells during development (B) and single colony screening (C). D) Primers used to sequence different portions of *KCNE1* and the corresponding barcode for subassembly. E) Histogram of barcode frequency during subassembly. Variant assignment was attempted for barcodes with >=500 reads (red line). Barcodes below this threshold were discarded.

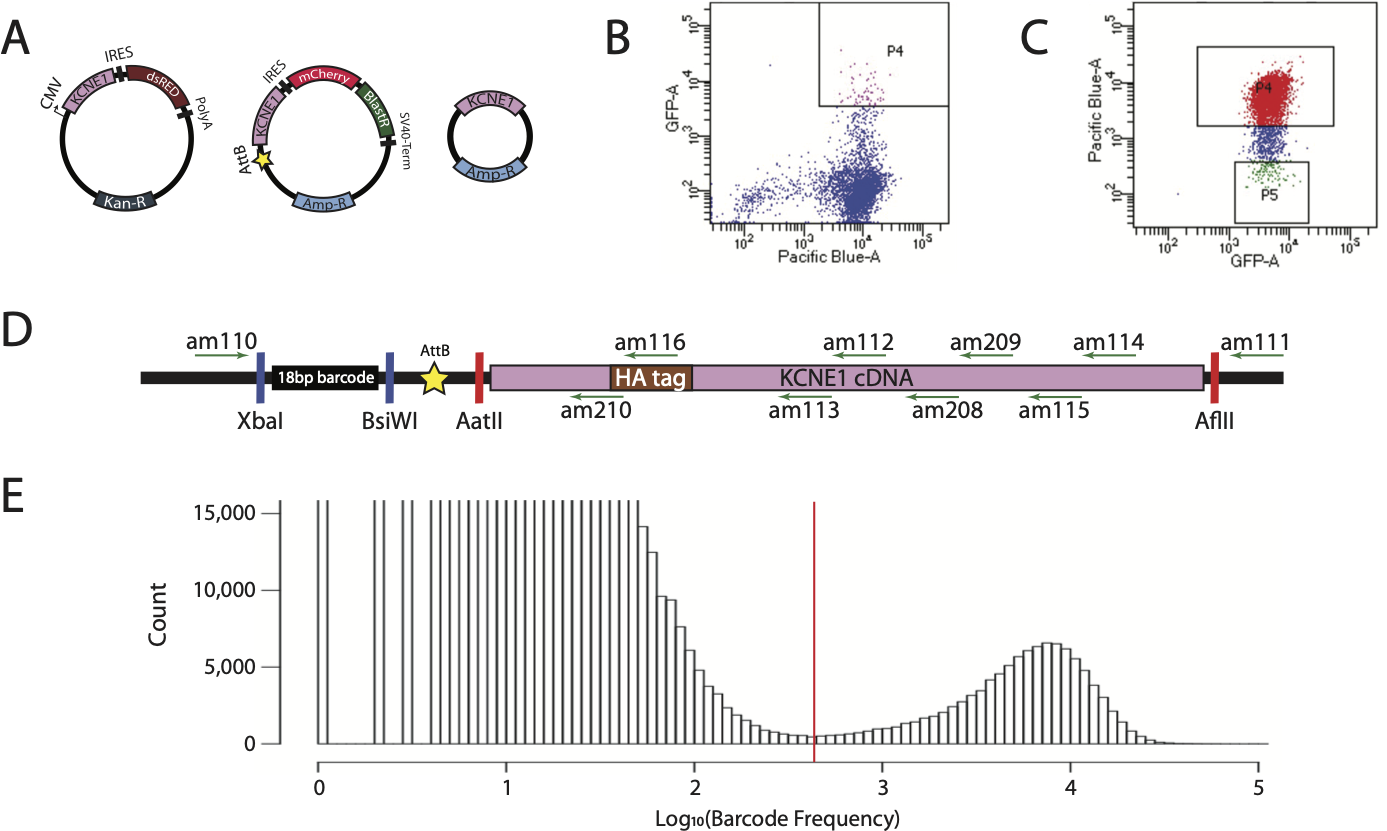

#### Figure S15: Flow cytometry gates for sorting cells based on cell surface KCNE1 expression

Flow cytometry gates for +KCNQ1 (A) and -KCNQ1 (B) trafficking experiments.

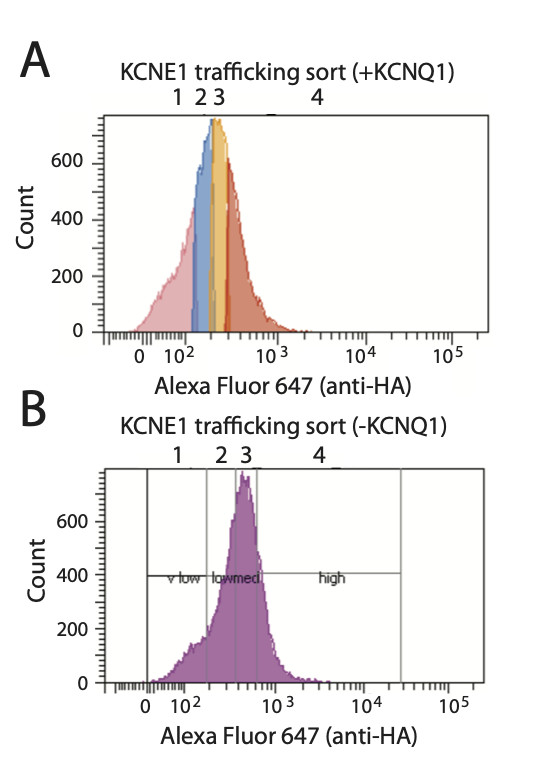

#### Figure S16: Convergence of the docking calculation of KCNE1 to KCNQ1 and calmodulin complex

The red dot represents the docked complex with the lowest binding energy (in Rosetta Energy Unit or REU; more negative values mean stronger binding). This complex was selected for structural analysis. Interface root-mean-square distance (r.m.s.d.) measures the average deviation of interface residues in all other docked models from the interface residues in the lowest-energy model. Interface residues are defined as residues in KCNE1 that are within 8Å from any KCNQ1 or calmodulin residues. The plot shows a typical “energy funnel” where binding energy becomes more negative as r.m.s.d. becomes smaller, suggesting the convergence of docking.

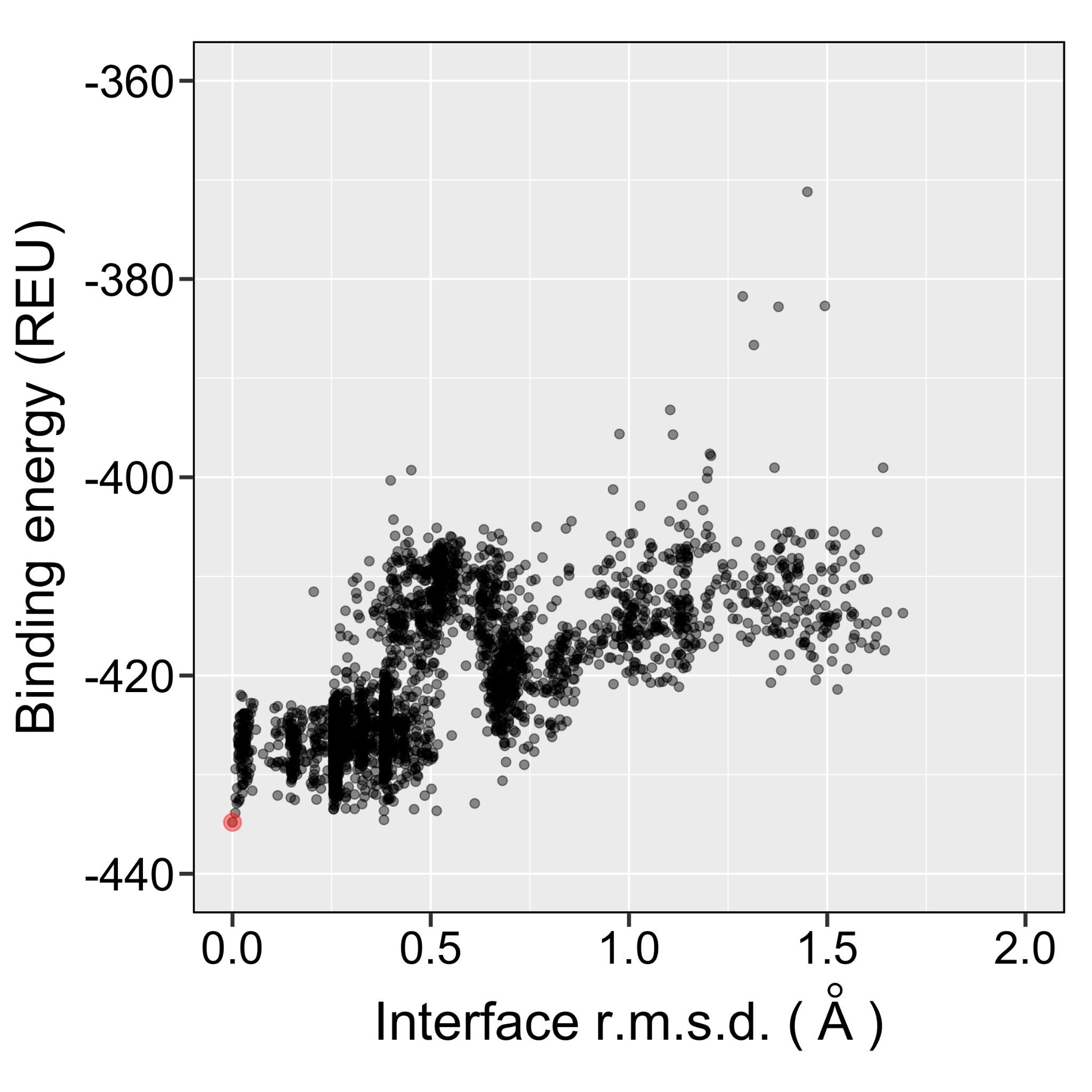

### Supplemental Tables

#### Table S1: Patch clamp data generated in this study

| **variant** | **Peak-20 (%)** | **Peak0 (%)** | **Peak20 (%)** | **Patch System** | **V1/2act (Mut-WT)** | **Peak summary** |
| --- | --- | --- | --- | --- | --- | --- |
| WT | 15.89 | 48.56 | 92.1 | HEK | 0 | 100 |
| I61X | 0 | 0.03 | 0.01 | HEK |  | 0.010857763 |
| G25V | 34.78 | 87.4 | 156.1 | HEK | -18.3 | 169.4896851 |
| G60D | 28.49 | 71.5 | 114.9 | HEK | -1.9 | 124.7557003 |
| D76N | 0.18 | 3.19 | 10.1 | HEK | -13.9 | 10.96634093 |
| G55X | 0.18 | 0.19 | 0.2 | HEK |  | 0.217155266 |
| F57Y | 2.54 | 14.86 | 48.78 | HEK | -1.9 | 52.96416938 |
| R98W | 4.73 | 15.26 | 31.72 | HEK | -12.8 | 34.44082519 |
| D85N | 32.3 | 117.22 | 215.59 | CHO | -20.09 | 234.082519 |
| F53X | 10.03 | 41.94 | 91.23 | CHO | 1.5 | 99.05537459 |
| L3P | 4.8 | 30.3 | 79.9 | CHO | 2.6 | 86.75352877 |
| E19D | 6.69 | 25.07 | 52.15 | HEK | -14.7 | 56.62323561 |
| D39G | 43.9 | 112.5 | 185.1 | HEK | -27.9 | 200.9771987 |
| E101V | 9.08 | 22.39 | 33.38 | HEK | -13.2 | 36.2432139 |
| T125M | 2.99 | 9.36 | 20.43 | HEK | -12.1 | 22.18241042 |
| P127T | 4.8 | 13.1 | 22.7 | CHO |  | 24.64712269 |

#### Table S2: Primers used in this study

| **Primer Name** | **Sequence** | **Description** |
| --- | --- | --- |
| ag289 | attacgtacgcttaagNNNNNNNNNNNNNNNNNNgacgtcctctagagcca | Barcode cloning forward |
| ag290 | tggctctagaggacgtc | Barcode cloning reverse |
| ag424 | taccagattatgcgccccgcagcggtgac | KCNE1 HA tag 34-35 forward |
| ag425 | catcgtaggggtaggacctgcgggccagg | KCNE1 HA tag 34-35 reverse |
| ag409 | attagcggccgcatgatcctgtctaacaccacagcg | KCNE1-HA move to pIRES2-dsRED2 |
| ag410 | attagcggccgctcatggggaaggcttcgtctc | KCNE1-HA move to pIRES2-dsRED2 |
| ag612 | attaggccatatggccatggccgcggcctc | KCNQ1 forward with SfiI |
| ag613 | attaggccatatggcctcaggacccctcatcggg | KCNQ1 reverse with SfiI |
| am5 | tctgcctcatcttcggcgtgctgtccacc | KCNQ1 S140G_Fwd |
| am6 | ggtggacagcacgccgaagatgaggcaga | KCNQ1 S140G_Rev |
| am10_PAGE | aatgatacggcgaccaccgagatctacacgccttgcgtctttccctacacgacgctcttccgatcttcttcgcccttagacaccat | NovaSeq Genomic DNA primers i5 |
| am11_PAGE | aatgatacggcgaccaccgagatctacactcctggtttctttccctacacgacgctcttccgatcttcttcgcccttagacaccat | NovaSeq Genomic DNA primers i5 |
| am12_PAGE | aatgatacggcgaccaccgagatctacaccttcaacctctttccctacacgacgctcttccgatcttcttcgcccttagacaccat | NovaSeq Genomic DNA primers i5 |
| am120_PAGE | caagcagaagacggcatacgagataggccgtggtgactggagttcagacgtgtgctcttccgatctcggcaattccggacgtacg | NovaSeq Genomic DNA primers i7 |
| am125 | aatgatacggcgaccaccgagatctacacacaacgcttctttccctacacgacgctcttccgatcttcttcgcccttagacaccat | NovaSeq Genomic DNA primers i5 |
| am126 | aatgatacggcgaccaccgagatctacacattccatatctttccctacacgacgctcttccgatcttcttcgcccttagacaccat | NovaSeq Genomic DNA primers i5 |
| am13_PAGE | aatgatacggcgaccaccgagatctacactcgctatgtctttccctacacgacgctcttccgatcttcttcgcccttagacaccat | NovaSeq Genomic DNA primers i5 |
| am14_PAGE | aatgatacggcgaccaccgagatctacacctatcgcatctttccctacacgacgctcttccgatcttcttcgcccttagacaccat | NovaSeq Genomic DNA primers i5 |
| am15_PAGE | aatgatacggcgaccaccgagatctacaccattagtgtctttccctacacgacgctcttccgatcttcttcgcccttagacaccat | NovaSeq Genomic DNA primers i5 |
| am20_PAGE | caagcagaagacggcatacgagataacgttaggtgactggagttcagacgtgtgctcttccgatctcggcaattccggacgtacg | NovaSeq Genomic DNA primers i7 |
| am21_PAGE | caagcagaagacggcatacgagatggtctattgtgactggagttcagacgtgtgctcttccgatctcggcaattccggacgtacg | NovaSeq Genomic DNA primers i7 |
| am22_PAGE | caagcagaagacggcatacgagataactcgccgtgactggagttcagacgtgtgctcttccgatctcggcaattccggacgtacg | NovaSeq Genomic DNA primers i7 |
| am23_PAGE | caagcagaagacggcatacgagatggagtgccgtgactggagttcagacgtgtgctcttccgatctcggcaattccggacgtacg | NovaSeq Genomic DNA primers i7 |
| am24_PAGE | caagcagaagacggcatacgagataagacattgtgactggagttcagacgtgtgctcttccgatctcggcaattccggacgtacg | NovaSeq Genomic DNA primers i7 |
| am25_PAGE | caagcagaagacggcatacgagattcgagccagtgactggagttcagacgtgtgctcttccgatctcggcaattccggacgtacg | NovaSeq Genomic DNA primers i7 |
| am30_PAGE | caagcagaagacggcatacgagatctagattggtgactggagttcagacgtgtgctcttccgatctcggcaattccggacgtacg | NovaSeq Genomic DNA primers i7 |
| am110-PAGE | caagcagaagacggcatacgagatggtaccgaccgtgactggagttcagacgtgtgctcttccgatct cggcaattccggacgtacg | NovaSeq sequencing primer for KCNE1 subassembly i7 |
| am111-PAGE | aatgatacggcgaccaccgagatctacacgtggtatctgacactctttccctacacgacgctcttccgatctccggtgcggccgccttaag | NovaSeq sequencing primer for KCNE1 subassembly i5 |
| am112-PAGE | aatgatacggcgaccaccgagatctacacgtggtatctgacactctttccctacacgacgctcttccgatctttcgagtgctccagcttct | NovaSeq sequencing primer for KCNE1 subassembly i5 |
| am113-PAGE | aatgatacggcgaccaccgagatctacacgtggtatctgacactctttccctacacgacgctcttccgatcttggagcggatgtagctcag | NovaSeq sequencing primer for KCNE1 subassembly i5 |
| am114-PAGE | aatgatacggcgaccaccgagatctacacgtggtatctgacactctttccctacacgacgctcttccgatctgtctcaggaaggtgtgtgt | NovaSeq sequencing primer for KCNE1 subassembly i5 |
| am115-PAGE | aatgatacggcgaccaccgagatctacacgtggtatctgacactctttccctacacgacgctcttccgatcttgggttgttctatggccag | NovaSeq sequencing primer for KCNE1 subassembly i5 |
| am116-PAGE | aatgatacggcgaccaccgagatctacacgtggtatctgacactctttccctacacgacgctcttccgatctaatctggtacatcgtaggggta | NovaSeq sequencing primer for KCNE1 subassembly i5 |
| am208 | aatgatacggcgaccaccgagatctacacgtggtatctgacactctttccctacacgacgctcttccgatctttgccaggcatcggactcga | NovaSeq sequencing primer for KCNE1 subassembly i5 |
| am209 | aatgatacggcgaccaccgagatctacacgtggtatctgacactctttccctacacgacgctcttccgatctctcttgccaggcatcggact | NovaSeq sequencing primer for KCNE1 subassembly i5 |
| am210 | aatgatacggcgaccaccgagatctacacgtggtatctgacactctttccctacacgacgctcttccgatcttaggggtaggacctgcgggc | NovaSeq sequencing primer for KCNE1 subassembly i5 |
| am180 | ccgcagcggtggcggcaagctgg | Quikchange D39G in KCNE1 |
| am182 | cgtctacatcgagtccaatgcctggcaagagaa | Quikchange D85N in KCNE1 |
| am183 | cccgggtcctggtgagctacaggtc | Quikchange E101V in KCNE1 |
| am179 | ccaagctgtggcaggatacagttcagcaggg | Quikchange E19D in KCNE1 |
| am185 | tcctcatggtactgggatagttcggcttcttcaccc | Quikchange F53X in KCNE1 |
| am191 | gggtgaagaagccgaactatcccagtaccatgagga | Quikchange F53X in KCNE1 |
| am187 | ggtactgggattcttcggcttctagaccctgggcatc | Quikchange F57X in KCNE1 |
| am192 | gatgcccagggtctagaagccgaagaatcccagtacc | Quikchange F57X in KCNE1 |
| am181 | ggtactgggattcttcggcttctataccctgggcatc | Quikchange F57Y in KCNE1 |
| am199 | gatgcccagggtatagaagccgaagaatcccagtacc | Quikchange F57Y in KCNE1 |
| am194 | agttcagcagggtgtcaacatgtcgggcc | Quikchange G25V in KCNE1 |
| am196 | ggcccgacatgttgacaccctgctgaact | Quikchange G25V in KCNE1 |
| am186 | ctcatggtactgggattcttctagttcttcaccctgggcatcatg | Quikchange G55X in KCNE1 |
| am198 | catgatgcccagggtgaagaactagaagaatcccagtaccatgag | Quikchange G55X in KCNE1 |
| am195 | ggcttcttcaccctggacatcatgctgagctac | Quikchange G60D in KCNE1 |
| am197 | gtagctcagcatgatgtccagggtgaagaagcc | Quikchange G60D in KCNE1 |
| am189 | gcttcttcaccctgggctagatgctgagctacatccg | Quikchange I61X in KCNE1 |
| am206 | cggatgtagctcagcatctagcccagggtgaagaagc | Quikchange I61X in KCNE1 |
| am178 | gcgacgtcatgatcccgtctaacaccacagc | Quikchange L3P in KCNE1 |
| am188 | cttcggcttcttcacctagggcatcatgctgagc | Quikchange L59X in KCNE1 |
| am205 | gctcagcatgatgccctaggtgaagaagccgaag | Quikchange L59X in KCNE1 |
| ag158 | gttcctgtagcggcggcgactctagatca | QuikChange mutate out NotI from pIRES2-dsRed2 |
| ag201 | tgtacaagtaaagcggcggcgactctagatcataa | Quikchange mutate out NotI from pIRES:GFP |
| bk196 | cttggcccgcgtccgccggtgag | QuikChange mutate out NotI in KCNQ1 |
| am184 | cacaccttcctgagatgaagccttccccatg | Quikchange T125M in KCNE1 |
| Maria's AdditionsA63:A63:I110 | | |
| ag579 | acctctacaaatgtggtatggctgattatgatcaggatccTTAGCCCTCCCACACATAACCA | Gibson AttB take 7 Rev primer |
| ag581 | gggccgccactccaccggcggcatggacgagctgtacaagGGTGGCAGCGGCGGTGGCAGTGGCGGTATGGCCAAGCCTTTGTCTCAAG | Gibson AttB take 7b (BlastR fusion) For primer |
| ag289_PAGE | attaCGTACGcttaagNNNNNNNNNNNNNNNNNNgacgtccTCTAGAgcca | Barcode for Zone 7 take 5 (PAGE purified) |
| ag290_PAGE | TGGCTCTAGAGGACGTC | Barcode for Zone 7 RC for making double stranded (PAGE purified) |
| ag1566 | GCCTGTCTAACACCACAGCGGT | KCNE1 I2C Jain F |
| ag1567 | ACATGACGTCGCGGCCG | KCNE1 I2C Jain R |
| bk331+bk332 |  | used to make pAG1296, which I used to clone KCNE1 sleeping beauty blue plasmid |
| Found in Primer Database mentioning KCNE1 | | |
| ag1568 | GTCAGCAGGGTGGCAACATGT | KCNE1 V21C Jain F |
| ag1569 | ATGTCTCCTGCCACAGCTTG | KCNE1 V21C Jain R |
| ag1570 | TCATGCTGAGCTACATCCGCTC | KCNE1 I61D Jain F WRONG |
| ag1571 | CGCCCAGGGTGAAGAAGC | KCNE1 I61D Jain R |
| ag1572 | ACATGCTGAGCTACATCCGCTC | KCNE1 I61D Jain F CORRECT |
| ag379 | TACCAGATTATGCGCAGGGTGG | WRONG Jain F KCNE1 HA tag 22-23 |
| ag380 | CATCGTAGGGGTACTGAACTGTCTC | WRONG Jain R KCNE1 HA tag 22-23 |
| ag381 | TACCAGATTATGCGCCCCGC | WRONG Jain F KCNE1 HA tag 34-35 |
| ag382 | CATCGTAGGGGTAGGACCTGC | WRONG Jain R KCNE1 HA tag 34-35 |
| ag409 | attaGCGGCCGCatgatcctgtctaacaccacagcg | KCNE1 into pIRES2 ag409+10-->400 bp-->NotI-->clone into pIRES2 |
| ag422 | TACCAGATTATGCGCAGGGTGGCAACATGTCGG | Jain F KCNE1 HA tag 22-23 correct |
| ag423 | CATCGTAGGGGTACTGAACTGTCTCCTGCCACAGC | Jain R KCNE1 HA tag 22-23 correct |
| ag424 | TACCAGATTATGCGCCCCGCAGCGGTGAC | Jain F KCNE1 HA tag 34-35 correct |
| ag425 | CATCGTAGGGGTAGGACCTGCGGGCCAGG | Jain R KCNE1 HA tag 34-35 correct |
| ag522 | attaGCGATCGCcttaagCAGGCTGAAGTTAGTAGCTCCGCTTCCggacccctcatcggggc | KCNQ1:KCNE1 zone 3R |
| ag523 | attaGCGGCCGCcttaagCAGGCTGGCGACGTGGAGGAGAACCCTGGACCTatgatcctgtctaacaccacagc | KCNQ1:KCNE1 zone 4F |
| ag524 | attaGCGGCCGCcttaagCAGGCTGAAGTTAGTAGCTCCGCTTCCggacccctcatcggggc | KCNQ1:KCNE1 Q1 only R |
| ag525 | attaGCGGCCGCaccggtTCATGGGGAAGGCTTCGTCTC | KCNQ1:KCNE1 full R |
| ag526 | cgatgaggggtccGGAAGCGGAGCTACTAACTTCAGCCTGcttaagCAGGCTGGCGACGTGG | KCNQ1:KCNE1 Z4GibsonF |
| ag527 | tgatcagttatctagatccggtggatcccggtGCGGCCGCaccggtTCATGGGGAAGGCTTCGTCTC | KCNQ1:KCNE1 Z4GibsonR |
| ag594 | gaggtcgacgatgtaggtcacggcattccggaGCGGCCGCGACGTCatgatcctgtctaacaccacagcg | Forward primer for cloning KCNE1 in minimalist |
| ag595 | aagctgcaataaacaagttaacaacaacaattgGCGATCGGCGGCCGCCTTAAGtcatggggaaggcttcgtct | Reverse primer for cloning KCNE1 in minimalist |
| ag596 | CTCTACGTCCTCATGGTACATGGATTCTTCGGCTTCTTCA | QC KCNE1 L51H |
| ag597 | CACACCTTCCTGAGACGAAGACTTCCCCATGA | QC KCNE1 P127T |
| ag609_PAGE | GTGACTGGAGTTCAGACGTGTGCTCTTCCGATCTacaacaattggcgatcggcg | Illumina Rev primer minimalist right of AsiSI (KCNE1 only) A |
| ag610_PAGE | GTGACTGGAGTTCAGACGTGTGCTCTTCCGATCTNNacaacaattggcgatcggcg | Illumina Rev primer minimalist right of AsiSI (KCNE1 only) B |
| ag611_PAGE | GTGACTGGAGTTCAGACGTGTGCTCTTCCGATCTNNNNacaacaattggcgatcggcg | Illumina Rev primer minimalist right of AsiSI (KCNE1 only) C |
| ag965 | attaGCGGCCGCatgatcctgtctaacaccacagc | NotI-KCNE1 forward primer for cloning into sleeping beauty vector |
| ag966 | attaGCGGCCGCtcatggggaaggcttcgtct | NotI-KCNE1 reverse primer for cloning into sleeping beauty vector |
| ag1417 | ACCAAGCTGTGGCAGAAGACAGTTCAGCAGG | KCNE1 E19K F |
| ag1418 | CCTGCTGAACTGTCTTCTGCCACAGCTTGGT | KCNE1 E19K R |
| ag1419 | GCGGTGACGGCGAGCTGGAGGCC | KCNE1 K41E F |
| ag1420 | GGCCTCCAGCTCGCCGTCACCGC | KCNE1 K41E R |
| ag1460 | GGCCCGGGTCCTGCTGAGCTACAGGTCG | KCNE1 QC E101L F |
| ag1461 | CGACCTGTAGCTCAGCAGGACCCGGGCC | KCNE1 QC E101L R |
| ag1462 | CAGGTCGTGCTATGTCGTTATAAACCATCTGGCCATAGAA | KCNE1 QC E110I F |
| ag1463 | TTCTATGGCCAGATGGTTTATAACGACATAGCACGACCTG | KCNE1 QC E110I R |
| ag1526 | AAGACAGTTCAGCAGGGTGG | KCNE1 E19K Jain F |
| ag1527 | CTGCCACAGCTTGGTCAGA | KCNE1 E19K Jain R |
| ag1528 | GAGCTGGAGGCCCTCTACG | KCNE1 K41E Jain F |
| ag1529 | GCCGTCACCGCTGCG | KCNE1 K41E Jain R |
| ag1530 | CTGAGCTACAGGTCGTGCTATGTC | KCNE1 E101L Jain F |
| ag1531 | CAGGACCCGGGCCTG | KCNE1 E101L Jain R |
| ag1532 | ATAAACCATCTGGCCATAGAACAACC | KCNE1 E110I Jain F |
| ag1533 | AACGACATAGCACGACCTGTAG | KCNE1 E110I Jain R |
| ag1818 | CCTGGAGAGCTACAGGTCGCTCTATGTCGTTGAAAACCAT | KCNE1 C106L quikchange primer |
| ag1819 | GGAGAGCTACAGGTCGTGCCGTGTCGTTGAAAACCATCTG | KCNE1 Y107R quikchange primer |
| ag1591 | AATGATACGGCGACCACCGAGATCTACACATGAGATCATACACTCTTTCCCTACACGACGCTCTTCCGATCTTCTTCGCCCTTAGACACCAT | NovaSeq i5 for amplifying from HEK |
| ag1592 | AATGATACGGCGACCACCGAGATCTACACGCAGAGCTGCACACTCTTTCCCTACACGACGCTCTTCCGATCTTCTTCGCCCTTAGACACCAT | NovaSeq i5 for amplifying from HEK |
| ag1593 | AATGATACGGCGACCACCGAGATCTACACTGTCGCTGGTACACTCTTTCCCTACACGACGCTCTTCCGATCTTCTTCGCCCTTAGACACCAT | NovaSeq i5 for amplifying from HEK |
| ag1594 | AATGATACGGCGACCACCGAGATCTACACCACTATCAACACACTCTTTCCCTACACGACGCTCTTCCGATCTTCTTCGCCCTTAGACACCAT | NovaSeq i5 for amplifying from HEK |
| ag1595 | AATGATACGGCGACCACCGAGATCTACACCTCTGCAGCGACACTCTTTCCCTACACGACGCTCTTCCGATCTTCTTCGCCCTTAGACACCAT | NovaSeq i5 for amplifying from HEK |
| ag1596 | AATGATACGGCGACCACCGAGATCTACACTCTCATGATAACACTCTTTCCCTACACGACGCTCTTCCGATCTTCTTCGCCCTTAGACACCAT | NovaSeq i5 for amplifying from HEK |
| ag1597 | AATGATACGGCGACCACCGAGATCTACACTATCTTGTAGACACTCTTTCCCTACACGACGCTCTTCCGATCTTCTTCGCCCTTAGACACCAT | NovaSeq i5 for amplifying from HEK |
| ag1598 | AATGATACGGCGACCACCGAGATCTACACCGCTCCACGAACACTCTTTCCCTACACGACGCTCTTCCGATCTTCTTCGCCCTTAGACACCAT | NovaSeq i5 for amplifying from HEK |
| ag1599 | AATGATACGGCGACCACCGAGATCTACACATTGCCGAGTACACTCTTTCCCTACACGACGCTCTTCCGATCTTCTTCGCCCTTAGACACCAT | NovaSeq i5 for amplifying from HEK |
| ag1600 | AATGATACGGCGACCACCGAGATCTACACGCCATTAGACACACTCTTTCCCTACACGACGCTCTTCCGATCTTCTTCGCCCTTAGACACCAT | NovaSeq i5 for amplifying from HEK |
| ag1601 | AATGATACGGCGACCACCGAGATCTACACAGCACATCCTACACTCTTTCCCTACACGACGCTCTTCCGATCTTCTTCGCCCTTAGACACCAT | NovaSeq i5 for amplifying from HEK |
| ag1602 | AATGATACGGCGACCACCGAGATCTACACGATGTGCTTCACACTCTTTCCCTACACGACGCTCTTCCGATCTTCTTCGCCCTTAGACACCAT | NovaSeq i5 for amplifying from HEK |
| ag1603 | CAAGCAGAAGACGGCATACGAGATGCCGACAAGAGTGACTGGAGTTCAGACGTGTGCTCTTCCGATCTcggcAattccggaCGTACG | NovaSeq i7 for amplifying from HEK |
| ag1604 | CAAGCAGAAGACGGCATACGAGATATTAGTGGAGGTGACTGGAGTTCAGACGTGTGCTCTTCCGATCTcggcAattccggaCGTACG | NovaSeq i7 for amplifying from HEK |
| ag1605 | CAAGCAGAAGACGGCATACGAGATCTGGCTTGCCGTGACTGGAGTTCAGACGTGTGCTCTTCCGATCTcggcAattccggaCGTACG | NovaSeq i7 for amplifying from HEK |
| ag1606 | CAAGCAGAAGACGGCATACGAGATTCAATCCATTGTGACTGGAGTTCAGACGTGTGCTCTTCCGATCTcggcAattccggaCGTACG | NovaSeq i7 for amplifying from HEK |
| ag1609 | CAAGCAGAAGACGGCATACGAGATCTGTTGGTCCGTGACTGGAGTTCAGACGTGTGCTCTTCCGATCTcggcAattccggaCGTACG | NovaSeq i7 for amplifying from HEK |
| ag1610 | CAAGCAGAAGACGGCATACGAGATTCACCAACTTGTGACTGGAGTTCAGACGTGTGCTCTTCCGATCTcggcAattccggaCGTACG | NovaSeq i7 for amplifying from HEK |
| ag1611 | CAAGCAGAAGACGGCATACGAGATGGTGCTATATGTGACTGGAGTTCAGACGTGTGCTCTTCCGATCTcggcAattccggaCGTACG | NovaSeq i7 for amplifying from HEK |
| ag1612 | CAAGCAGAAGACGGCATACGAGATAACATCGCGCGTGACTGGAGTTCAGACGTGTGCTCTTCCGATCTcggcAattccggaCGTACG | NovaSeq i7 for amplifying from HEK |
| ag1613 | CAAGCAGAAGACGGCATACGAGATGGTGGACGTGGTGACTGGAGTTCAGACGTGTGCTCTTCCGATCTcggcAattccggaCGTACG | NovaSeq i7 for amplifying from HEK |
| ag1614 | CAAGCAGAAGACGGCATACGAGATAACAAGTACAGTGACTGGAGTTCAGACGTGTGCTCTTCCGATCTcggcAattccggaCGTACG | NovaSeq i7 for amplifying from HEK |
| ag1615 | CAAGCAGAAGACGGCATACGAGATACCGTTACAAGTGACTGGAGTTCAGACGTGTGCTCTTCCGATCTcggcAattccggaCGTACG | NovaSeq i7 for amplifying from HEK |
| ag1616 | CAAGCAGAAGACGGCATACGAGATGTTACCGTGGGTGACTGGAGTTCAGACGTGTGCTCTTCCGATCTcggcAattccggaCGTACG | NovaSeq i7 for amplifying from HEK |
| ag1426 | CAAGCAGAAGACGGCATACGAGATTGACGACCACGTGACTGGAGTTCAGACGTGTGCTCTTCCGATCTTCTTCGCCCTTAGACACCAT | NovaSeq Post-cell barcode i7 primer index for Vantage GTGGTCGTCA, use with ag1436 |
| ag1427 | CAAGCAGAAGACGGCATACGAGATCAGTAGTTGTGTGACTGGAGTTCAGACGTGTGCTCTTCCGATCTTCTTCGCCCTTAGACACCAT | NovaSeq Post-cell barcode i7 primer index for Vantage ACAACTACTG, use with ag1437 |
| ag1428 | CAAGCAGAAGACGGCATACGAGATCTCTCGAGCCGTGACTGGAGTTCAGACGTGTGCTCTTCCGATCTTCTTCGCCCTTAGACACCAT | NovaSeq Post-cell barcode i7 primer index for Vantage GGCTCGAGAG, use with ag1438 |
| ag1436 | AATGATACGGCGACCACCGAGATCTACACACTGTTGTGAACACTCTTTCCCTACACGACGCTCTTCCGATCTNNNNNNNNNNcggcAattccggaCGTACG | NovaSeq Post-cell barcode i5 primer index for Vantage TCACAACAGT, use with ag1426, left of barcode with UMI10 |
| ag1437 | AATGATACGGCGACCACCGAGATCTACACGTCACCACAGACACTCTTTCCCTACACGACGCTCTTCCGATCTNNNNNNNNNNcggcAattccggaCGTACG | NovaSeq Post-cell barcode i5 primer index for Vantage CTGTGGTGAC, use with ag1427, left of barcode with UMI10 |
| ag1438 | AATGATACGGCGACCACCGAGATCTACACCGCTCAGTTCACACTCTTTCCCTACACGACGCTCTTCCGATCTNNNNNNNNNNcggcAattccggaCGTACG | NovaSeq Post-cell barcode i5 primer index for Vantage GAACTGAGCG, use with ag1428, left of barcode with UMI10 |
| ag1426 | CAAGCAGAAGACGGCATACGAGATTGACGACCACGTGACTGGAGTTCAGACGTGTGCTCTTCCGATCTTCTTCGCCCTTAGACACCAT | NovaSeq Post-cell barcode i7 primer index for Vantage GTGGTCGTCA, use with ag1436 |
| ag1427 | CAAGCAGAAGACGGCATACGAGATCAGTAGTTGTGTGACTGGAGTTCAGACGTGTGCTCTTCCGATCTTCTTCGCCCTTAGACACCAT | NovaSeq Post-cell barcode i7 primer index for Vantage ACAACTACTG, use with ag1437 |
| ag1428 | CAAGCAGAAGACGGCATACGAGATCTCTCGAGCCGTGACTGGAGTTCAGACGTGTGCTCTTCCGATCTTCTTCGCCCTTAGACACCAT | NovaSeq Post-cell barcode i7 primer index for Vantage GGCTCGAGAG, use with ag1438 |
| ag1429 | CAAGCAGAAGACGGCATACGAGATTCTCTAGATTGTGACTGGAGTTCAGACGTGTGCTCTTCCGATCTTCTTCGCCCTTAGACACCAT | NovaSeq Post-cell barcode i7 primer index for Vantage AATCTAGAGA, use with ag1439 |
| ag1430 | CAAGCAGAAGACGGCATACGAGATGGTAACGCAGGTGACTGGAGTTCAGACGTGTGCTCTTCCGATCTTCTTCGCCCTTAGACACCAT | NovaSeq Post-cell barcode i7 primer index for Vantage CTGCGTTACC, use with ag1440 |
| ag1431 | CAAGCAGAAGACGGCATACGAGATAACGGTATGAGTGACTGGAGTTCAGACGTGTGCTCTTCCGATCTTCTTCGCCCTTAGACACCAT | NovaSeq Post-cell barcode i7 primer index for Vantage TCATACCGTT, use with ag1441 |
| ag1432 | CAAGCAGAAGACGGCATACGAGATTCGACTTAAGGTGACTGGAGTTCAGACGTGTGCTCTTCCGATCTTCTTCGCCCTTAGACACCAT | NovaSeq Post-cell barcode i7 primer index for Vantage CTTAAGTCGA, use with ag1442 |
| ag1433 | CAAGCAGAAGACGGCATACGAGATCTAGTCCGGAGTGACTGGAGTTCAGACGTGTGCTCTTCCGATCTTCTTCGCCCTTAGACACCAT | NovaSeq Post-cell barcode i7 primer index for Vantage TCCGGACTAG, use with ag1443 |
| ag1434 | CAAGCAGAAGACGGCATACGAGATGAGCAAGGCCGTGACTGGAGTTCAGACGTGTGCTCTTCCGATCTTCTTCGCCCTTAGACACCAT | NovaSeq Post-cell barcode i7 primer index for Vantage GGCCTTGCTC, use with ag1444 |
| ag1435 | CAAGCAGAAGACGGCATACGAGATAGATGGAATTGTGACTGGAGTTCAGACGTGTGCTCTTCCGATCTTCTTCGCCCTTAGACACCAT | NovaSeq Post-cell barcode i7 primer index for Vantage AATTCCATCT, use with ag1445 |
| ag1436 | AATGATACGGCGACCACCGAGATCTACACACTGTTGTGAACACTCTTTCCCTACACGACGCTCTTCCGATCTNNNNNNNNNNcggcAattccggaCGTACG | NovaSeq Post-cell barcode i5 primer index for Vantage TCACAACAGT, use with ag1426, left of barcode with UMI10 |
| ag1437 | AATGATACGGCGACCACCGAGATCTACACGTCACCACAGACACTCTTTCCCTACACGACGCTCTTCCGATCTNNNNNNNNNNcggcAattccggaCGTACG | NovaSeq Post-cell barcode i5 primer index for Vantage CTGTGGTGAC, use with ag1427, left of barcode with UMI10 |
| ag1438 | AATGATACGGCGACCACCGAGATCTACACCGCTCAGTTCACACTCTTTCCCTACACGACGCTCTTCCGATCTNNNNNNNNNNcggcAattccggaCGTACG | NovaSeq Post-cell barcode i5 primer index for Vantage GAACTGAGCG, use with ag1428, left of barcode with UMI10 |
| ag1439 | AATGATACGGCGACCACCGAGATCTACACTATCTGACCTACACTCTTTCCCTACACGACGCTCTTCCGATCTNNNNNNNNNNcggcAattccggaCGTACG | NovaSeq Post-cell barcode i5 primer index for Vantage AGGTCAGATA, use with ag1429, left of barcode with UMI10 |
| ag1440 | AATGATACGGCGACCACCGAGATCTACACCCGTGGCCTTACACTCTTTCCCTACACGACGCTCTTCCGATCTNNNNNNNNNNcggcAattccggaCGTACG | NovaSeq Post-cell barcode i5 primer index for Vantage AAGGCCACGG, use with ag1430, left of barcode with UMI10 |
| ag1441 | AATGATACGGCGACCACCGAGATCTACACTTACAATTCCACACTCTTTCCCTACACGACGCTCTTCCGATCTNNNNNNNNNNcggcAattccggaCGTACG | NovaSeq Post-cell barcode i5 primer index for Vantage GGAATTGTAA, use with ag1431, left of barcode with UMI10 |
| ag1442 | AATGATACGGCGACCACCGAGATCTACACAGTGGTCAGGACACTCTTTCCCTACACGACGCTCTTCCGATCTNNNNNNNNNNcggcAattccggaCGTACG | NovaSeq Post-cell barcode i5 primer index for Vantage CCTGACCACT, use with ag1432, left of barcode with UMI10 |
| ag1443 | AATGATACGGCGACCACCGAGATCTACACGACAACTGAAACACTCTTTCCCTACACGACGCTCTTCCGATCTNNNNNNNNNNcggcAattccggaCGTACG | NovaSeq Post-cell barcode i5 primer index for Vantage TTCAGTTGTC, use with ag1433, left of barcode with UMI10 |
| ag1444 | AATGATACGGCGACCACCGAGATCTACACACGGAATGCGACACTCTTTCCCTACACGACGCTCTTCCGATCTNNNNNNNNNNcggcAattccggaCGTACG | NovaSeq Post-cell barcode i5 primer index for Vantage CGCATTCCGT, use with ag1434, left of barcode with UMI10 |
| ag1445 | AATGATACGGCGACCACCGAGATCTACACGTAAGGCATAACACTCTTTCCCTACACGACGCTCTTCCGATCTNNNNNNNNNNcggcAattccggaCGTACG | NovaSeq Post-cell barcode i5 primer index for Vantage TATGCCTTAC, use with ag1435, left of barcode with UMI10 |
| ag1581 | CAAGCAGAAGACGGCATACGAGATCAACATTCAAGTGACTGGAGTTCAGACGTGTGCTCTTCCGATCTTCTTCGCCCTTAGACACCAT | NovaSeq Post-cell barcode i7 primer index for Vantage TTGAATGTTG, use with ag1583 |
| ag1582 | CAAGCAGAAGACGGCATACGAGATTAACTTAGCGGTGACTGGAGTTCAGACGTGTGCTCTTCCGATCTTCTTCGCCCTTAGACACCAT | NovaSeq Post-cell barcode i7 primer index for Vantage CGCTAAGTTA, use with ag1584 |
| ag1583 | AATGATACGGCGACCACCGAGATCTACACCCTCGCAACCACACTCTTTCCCTACACGACGCTCTTCCGATCTNNNNNNNNNNcggcAattccggaCGTACG | NovaSeq Post-cell barcode i5 primer index for Vantage GGTTGCGAGG, use with ag1581, left of barcode with UMI10 |
| ag1584 | AATGATACGGCGACCACCGAGATCTACACAGGTGCGTAAACACTCTTTCCCTACACGACGCTCTTCCGATCTNNNNNNNNNNcggcAattccggaCGTACG | NovaSeq Post-cell barcode i5 primer index for Vantage TTACGCACCT, use with ag1582, left of barcode with UMI10 |

#### Table S3: List of previously developed computational metrics compared to MAVE scores in this study

| **Name of Metric** | **Reference** | **Year published** |
| --- | --- | --- |
| SIFT_score | Ng and Henikoff | 2003 |
| Polyphen2_HVAR_score | Adzhubei et al. | 2010 |
| LRT_score | Chun and Fay | 2009 |
| MutationTaster_score (v2) | Schwarz et al. | 2014 |
| MutationAssessor_score | Reva et al. | 2011 |
| FATHMM_score (weighted) | Shihab et al. | 2013 |
| PROVEAN_score | Choi and Chan | 2015 |
| VEST4_score | Carter et al. | 2013 |
| MetaSVM_score | Dong et al. | 2015 |
| MetaLR_score | Dong et al. | 2015 |
| REVEL_score | Ioannidis et al. | 2016 |
| MutPred_score (v1.2) | Li et al. | 2009 |
| MVP_score | Qi et al. | 2018 |
| MPC_score | Samocha et al. | 2017 |
| CADD_raw | Kircher et al. | 2014 |
| GERP++_RS | Davydov et al. | 2010 |
| phyloP100way_vertebrate | Pollard et al. | 2010 |
| phastCons100way_vertebrate | Siepel et al. | 2005 |
